# Multiplexed imaging mass cytometry reveals distinct tumor-immune microenvironments linked to immunotherapy responses in melanoma

**DOI:** 10.1101/2022.01.15.476480

**Authors:** Xu Xiao, Qian Guo, Chuanliang Cui, Yating Lin, Lei Zhang, Xin Ding, Qiyuan Li, Minshu Wang, Wenxian Yang, Yan Kong, Rongshan Yu

## Abstract

Single-cell technologies have enabled extensive analysis of complex immune composition, phenotype and interactions within tumor, which is crucial in understanding the mechanisms behind cancer progression and treatment resistance. Unfortunately, the knowledge on cell phenotypes and their spatial interactions at present has only limited utilization in guiding pathological stratification on patients based on their immune microenvironments for better clinical decisions. Here we used imaging mass cytometry (IMC) to simultaneously quantify 35 proteins in a spatially resolved manner on tumor tissues from melanoma patients receiving anti-programmed cell death-1 (anti-PD-1) therapy. Unbiased single-cell analysis revealed highly dynamic tumor-immune microenvironments that are characterized with variable tumor and immune cell phenotypes and their organizations across and within melanomas, and identified distinct archetypes of melanoma microenvironments that are associated with benefit from anti-PD-1 therapy based on high-dimensional multicellular features. Our results demonstrate the utility of multiplex proteomic imaging technologies in studying complex molecular events in a spatially resolved manner for the development of new strategies for patient stratification and treatment outcome prediction.

---

Advanced melanoma had poor prognosis with a 5-year survival rate lower than 10%^1^. Immune checkpoint inhibitors (ICIs) targeting PD-1 and CTLA-4 have shown improved survival in advanced melanoma patients^1–4^, but potent and durable response only presented in a subset of patients. To date, no single biomarker has been sufficient for patient stratification, presumably due to the complex immune response to cancer driven by both inter-and intra-patient cellular heterogeneities in tumor microenvironments (TMEs). Indeed, with deeper knowledge on the mechanisms of immune checkpoint blockade (ICB) based immunotherapy developed from recent clinical and preclinical studies, it is now recognized that ICI efficacy is driven by multifaceted interactions among a large diversity of cell lineages at both localized and systemic levels^5–13^, thus defying the concept of patient stratification based solely on biomarkers that capture only limited dimensions of these intricate interactions.

In this study, we used IMC to explore the composition and spatial arrangements of different immune and stromal cells in the vicinity of cancer cells in baseline tumor samples from 26 advanced melanoma patients treated with anti-PD-1 monoclonal antibody at Peking University Cancer Hospital (PUCH), Beijing, China. Using single-cell analysis on high-dimensional mass cytometry images, we quantified inter-and intra-tumor heterogeneities in a spatially resolved manner and identified important cellular features to classify melanoma into distinct archetypes linked to immunotherapy outcome.

## Results

### Global characteristics of cell compositions in melanoma TME

We used a customized IMC panel of 35 antibodies targeting markers of tumor, immune, and stromal cell phenotypes, immuno-regulatory proteins, and proteins providing insights into cell activation, proliferation, and metabolism status (Supplementary Table 2) on baseline tissue samples from 26 melanoma patients treated with anti-PD-1 (Fig. 1a, Supplementary Table 1). Regions of interest (ROIs) were randomly selected for each sample from core tumor (CT) and invasive margin (IM) regions based on hematoxylin and eosin (HE)-stained serial tissue section inspected by a professional pathologist. After quality control by manual inspections, 158 IMC images (59 from the CT: 34 responders, 25 nonresponders, 99 from the IM: 58 responders, 41 nonresponders) were further analyzed.

**Figure 1:**
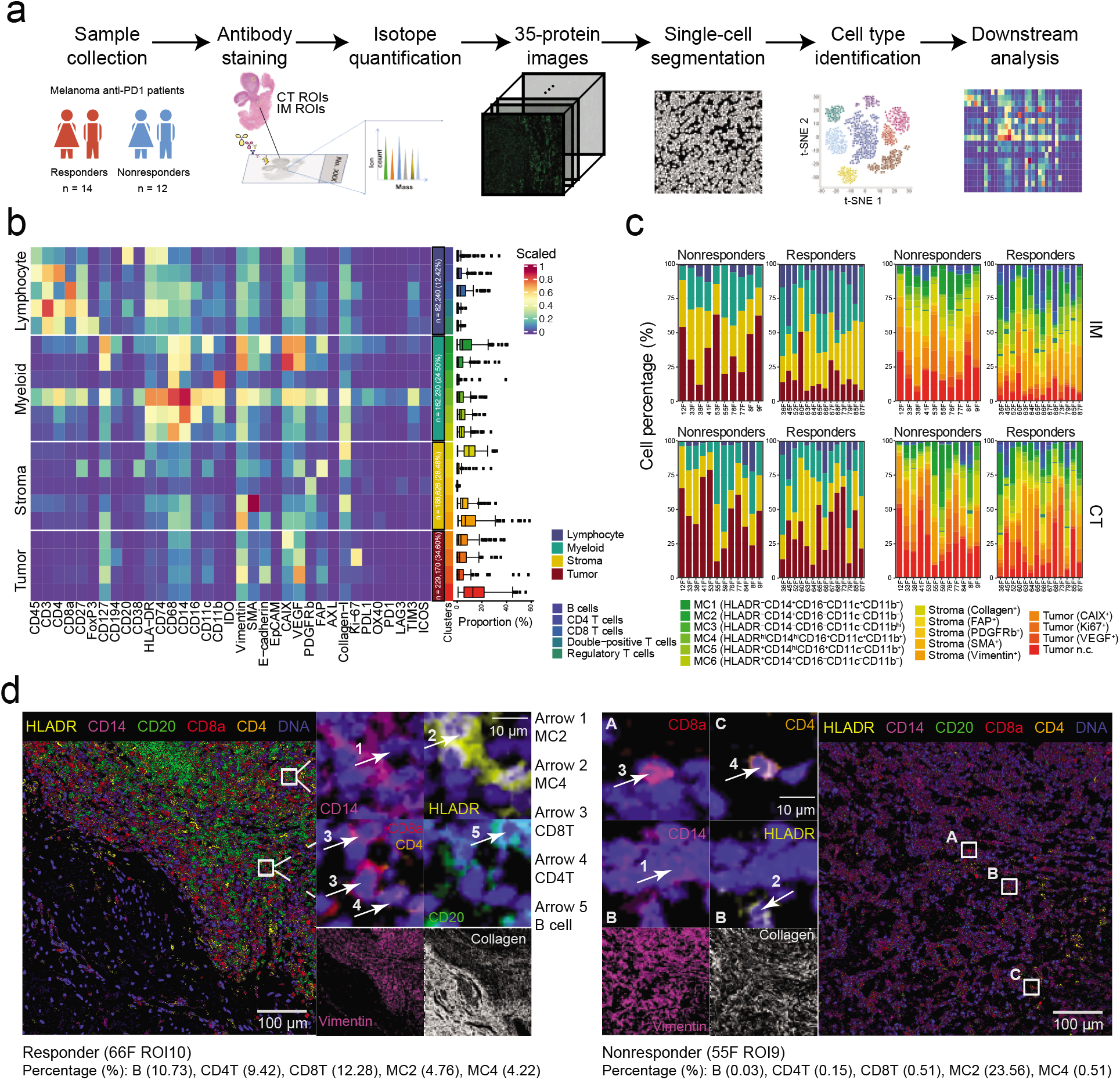
Overview of the study of melanoma patients using IMC and characteristics of cell composition in TMEs of melanoma. (a) Workflow of IMC images acquisition from melanoma patients and data analyses. (b) Heatmap of mean values of scaled protein expression per cell type identified by unsupervised clustering (FlowSOM and Phenograph) for a total of 662,266 single cells. The boxplots on the right depict the proportion of each cluster per IMC image. (c) Stack bars showing averaged cell percentage in images in the IM (top) and CT (bottom) from responders and nonresponders, colored by four main cell types (left) and 20 subtypes (right). (d) Representative multichannel IMC images from one responder and one nonresponder. Vimentin (magenta) and collagen I (white) were used to portrait the structure of tissue.

In total, 662,266 cells were clustered into 20 different cell subtypes using FlowSOM^14^ and Phenograph^15^ (Methods), which were further grouped into four major cell types including lympho-cytes, myeloid derived monocytes, stromal cells and tumor cells (Fig. 1b, Supplementary Fig. S1a-b). The lymphocytes included five different subtypes, namely, CD4 T cell (CD3^+^CD4^+^), CD8 T cell (CD3^+^CD8^+^), double-positive T cell (DPT; CD4^+^CD8^+^), T regulatory cell (Treg; CD4^+^FOXP3^+^) and B cell (CD19^+^) identified by their canonical cell markers. Myeloid derived monocytes (MC1-MC6) were identified by CD14 and CD16, which can be classified into two categories based on their MHC Class II molecule (HLA-DR) expression. The first category included three subtypes characterized with highly elevated HLA-DR expression (MC4-MC6), indicative of their potential role as antigen presenting cells (APC) within TME. Among them, subtype MC4 was further characterized with elevated dendritic cell marker CD11c and MC6 with elevated macrophage marker CD68. The second category was comprised of HLA-DR*^−^* subtypes with elevated expression of exhaustion markers CAIX and VEGF (MC2, MC3) or indoleamine 2,3-dioxygenase 1 (IDO-1; MC1), representing their potential immune suppressive roles as myeloid-derived suppressor cells (MDSCs). Stromal cells consisted of 5 subtypes denoted as S1 to S5 for Collagen^+^, FAP^+^, PDGFRb^+^, SMA^+^, and Vimentin^+^ cells, respectively, and tumor cells included 4 subtypes denoted as T1 to T4 for CAIX^+^, Ki67^+^, VEGF^+^, and a non-classified subtype (n.c.) that did not show elevated expression on any markers from the defined panel, respectively.

All major cell types and subtypes were observed in all patients but with significant variation in cell compositions among patients and different tumor regions (Fig. 1c). Overall, the IM demonstrated more diversified cell type compositions as indicated by higher Shannon entropy (Methods) than the CT for both responders and nonresponders (Supplementary Fig. S1c). Furthermore, Shannon entropy analysis indicated more diversified cell type compositions in responders than in non-responders in the IM, but not in the CT. Two IMC images to exemplify TMEs with typical immune cells in a responder and a nonresponder are shown in Fig. 1d.

### Cell phenotype proportions differentiate TMEs of responders and nonresponders

Examination of abundances of individual cell clusters from different TMEs revealed significantly different cell compositions in TMEs from responders and nonresponders. The percentages of lymphocytes were significantly higher in responders than in nonresponders in the IM but not CT (Fig. 2a-b). Similar trend was observed in all 5 lymphocyte subtypes, indicating the important role of the IM in identifying TMEs that would respond to immunotherapy. Interestingly, despite the well-established immunosuppressive role of Treg, significantly elevated Treg densities were observed in the IM of responders compared to nonresponders, which were possibly recruited to the site for maintaining immunological unresponsiveness to self-antigens and suppressing excessive immune responses detrimental to the host. As a result, high abundance of Treg could indicate the presence of highly immunogenic tumor associated antigens that would be able to induce a T cell mediated immune response after ICB for cancer rejection. For myeloid cells, we identified that HLA-DR^+^ myeloid cells MC4 were significantly more abundant in responders, while HLA-DR*^−^* myeloid cells MC2 were significantly enriched in nonresponders and the difference can be observed in both the IM and CT. We also found that tumor cells with hypoxia signals (CAIX^+^) were significantly enriched in the IM from nonresponders compared to responders, but this difference was not observed in the CT (Supplementary Fig. S2). No significant differences in other cell type abundances were observed.

**Figure 2:**
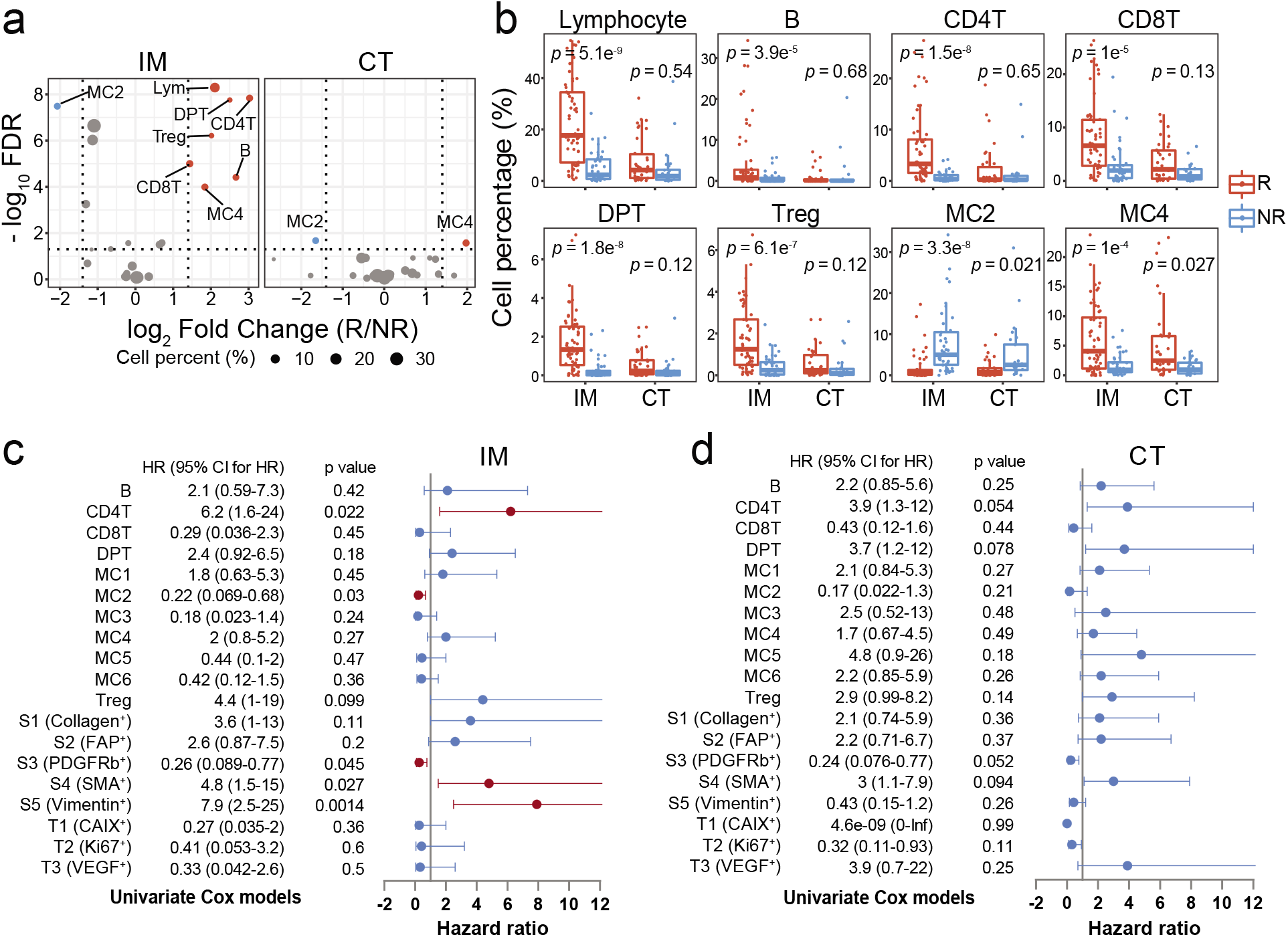
The prognostic impact of cell phenotypes in TME. (a) Volcano plots showing differential testing of cell abundance in the IM (left) and CT (right) between responders (R) and nonresponders (NR). The color of the nodes represents significant higher abundance (red) and lower abundance (blue) of cell type in responders. The size of the nodes displays the percentage of cell type. (b) Boxplots showing the proportion of cell type in ROIs from responders (R, red) and nonresponders (NR, blue). Each boxplot is shown with the median (the center line), interquartile range (IQR), and 1.5 times the IQR (whiskers), with outliers exceeding 1.5 times the IQR. Points in the boxplot represent the cell percentage of each IMC image (IM: n = 99, CT: n = 59). Comparisons were performed using Wilcoxon rank sum test and adjusted with Benjamini-Hochberg method. (c-d) Forest plots showing hazard ratios (nodes) and 95% confidence intervals (horizontal lines) of PFS for each cell type in the IM (c) and CT (d) by univariate Cox models adjusted for age. The red nodes represent the significant factor with *p* value *<* 0.05.

Cox regression analysis further revealed that the abundances of several cell types in the IM were associated with immunotherapy outcome. In the IM, CD4 T, SMA^+^ stromal cells S4, and Vimentin^+^ stromal cells S5 were associated with better outcome, whereas HLA-DR*^−^* myeloid cell MC2 and PDGFRb^+^ stromal cell S3 were indicative of poor outcome after adjusted for age (Fig. 2c). None of the identified cell phenotypes in the CT was prognostic (Fig. 2d).

### Characteristics of checkpoint expressions in TME

We next investigated the expressions of checkpoint molecules on different cell subtypes within TMEs to see if the compositions of any cell subtypes are associated with outcome to ICI treatments. Overall, PD-L1 was expressed on a broad class of tumor and stromal cells within TMEs from both responders and nonresponders, with MC4 having the highest average of PD-L1^+^ proportion (Supplementary Fig. S3a). However, none of their relative abundances, i.e., the percentages of PD-L1+ cells among the corresponding cell subtypes, was associated with response. Instead, significantly higher relative densities of PD-1^+^ CD4 T and CD8 T were observed in the IM of responders than that of nonresponders (Supplementary Fig. S3b), which is consistent with previous results that the fraction of exhausted cytotoxic T lymphocytes expressing high levels of CTLA-4 and PD-1 strongly correlates with responses to anti-PD-1 in human melanoma^16^.

In addition to PD-1, we also observed increased relative abundances of CD27 and TIM-3 positive cells among a broad class of lymphocyte and myeloid subtypes in the IM of responders (Supplementary Fig. S3c-d). CD27 is typically upregulated in the memory phenotypes of T cells upon exposure to stimulation^17^. In addition to their assumed roles in local immunity control, memory CD8^+^ T cells can further orchestrate the generation of systemic anti-tumor immunity by triggering antigen spreading through DC^18^, and the presence of resident memory T cells is associated with durable responses to immunotherapy in metastatic melanoma^19^. TIM-3 is a checkpoint receptor expressed on immune cells from TME including interferon (IFN)-*γ*-producing T cells and other leukocytes as well including DC and natural killer (NK) cells^20^. Although elevated expression of TIM-3 within TME was typically associated with T-cell exhaustion, a recent study showed that lack of TIM-3 expression of T cells may indicate a specific dysfunction status of T cells from ICB-refractory TMEs despite a brisk T cell infiltrate^21^. In addition, a preclinical study using a murine model of head and neck cancer showed that the suppressive activity of TIM-3 can be reversed by IFN-*γ* secreted by CD8^+^ T cells upon PD-1 blockade^22^. These observations, together with the results described earlier, suggest the potential clinical utilization of predicting outcome to PD-1 based ICB therapy based on sigatures of activated or previously activated antigen-experienced lymphocytes in the IM of tumor.

### Spatial analysis reveals heterogeneous cell-cell interactions in melanoma TME

We performed regional correlation analysis to investigate the potential spatial co-occurrence patterns of different cells across all images, and permutation-test-based neighbourhood analysis^23^ to identify statistically significant interaction or avoidance between pairs of cell types in melanoma (Methods, Fig. 3a-b, examples of cell-cell interaction are shown in Fig. 3c-g). Notably, subtypes of lymphocytes (CD4 T, CD8 T, DPT, B cells and Treg) tended to form dense compartments with strong cognate interactions and their proportions were highly correlated across images in responders (Fig. 3a, highlighted area 1, Fig. 3c-d). In nonresponders, although the positive correlations between different lymphocyte subtypes were still maintained, co-locations of these lymphocytes, particularly between CD4 T and other T cell subtypes, were observed in fewer ROIs (Fig. 3b, highlighted area 1; Supplementary Fig. S4a), indicative of a more diffused distribution of lymphocytes in these TMEs. We also observed highly different interaction patterns of HLA-DR^+^ and HLA-DR*^−^* myeloid cells with lymphocytes. Strong cognate interaction between the HLA-DR^+^CD11c^+^ myeloid cells (MC4) and lymphocytes can be observed in responders (Fig. 3a, highlighted area 2, Fig. 3e) and, to a lesser extent, in nonresponders as well (Fig. 3b, highlighted area 2; Supplementary Fig. S4a). In contrast, none of the HLA-DR*^−^* myeloid cells (MC1-MC3) interacted with lymphocytes. Instead, consistent with their immune suppressive functions, evidences of avoidance between these myeloid cells and lymphocytes from both abundance correlations and neighbourhood analysis can be observed in both responders and nonresponders (Fig. 3a, 3b, highlighted area 3, Supplementary Fig. S4b). Significant proximate interaction between SMA^+^ stromal cells, which are primarily vascular smooth muscle cells that surround lymphatic vessels or blood vessels, and a broad class of immune cells were observed in most ROIs from both responders and nonresponders (Fig. 3a, 3b, highlighted area 4, Fig. 3f-g), indicative of the significant role of lymphovascular structures in maintaining the immune cell populations in TME.

**Figure 3:**
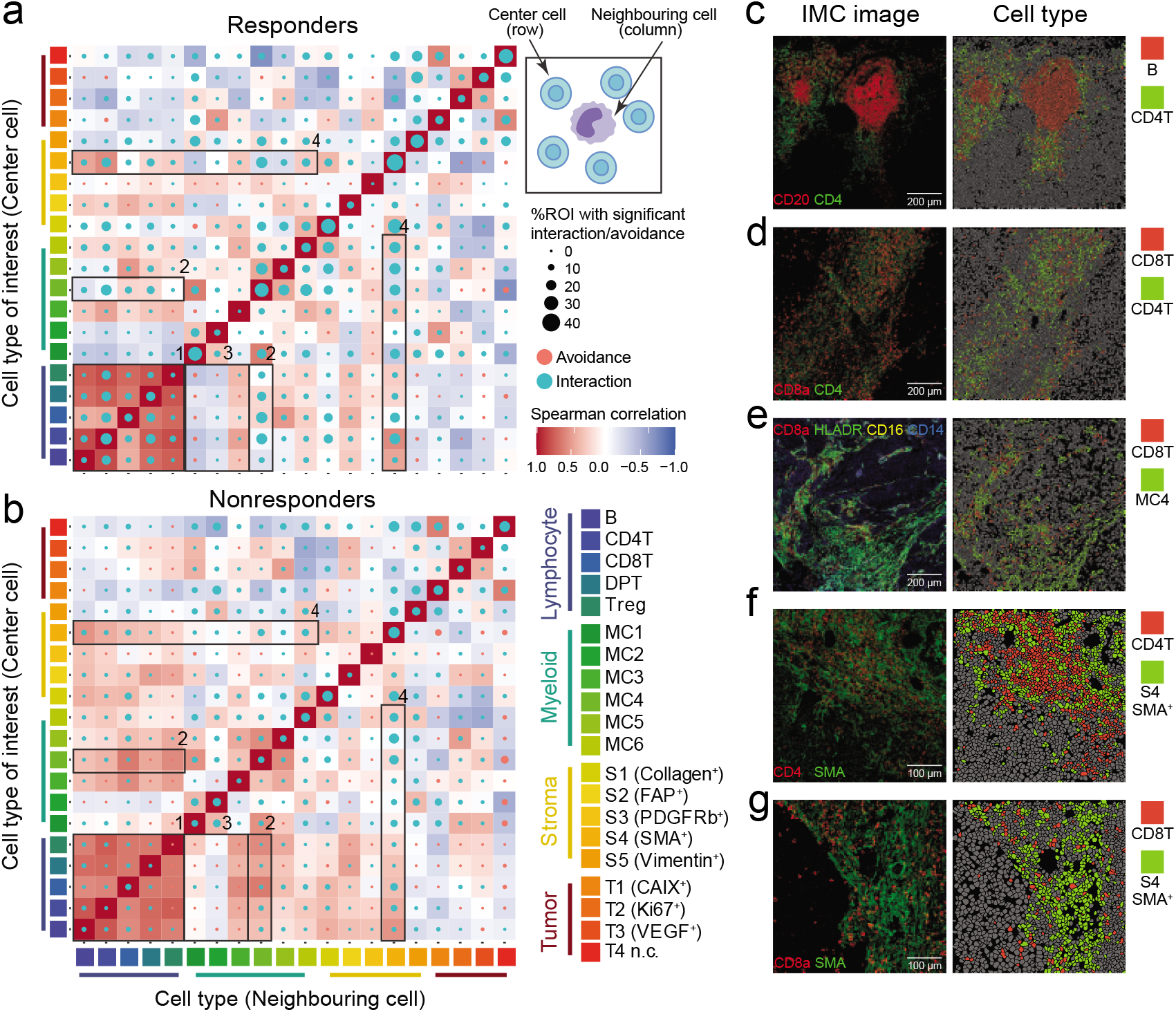
Spatial analysis among cell phenotypes. (a-b) Circles indicating patterns of cell-cell interactions (green) or avoidance (red) for responders (a) and nonresponders (b). The circle size showing the percentage of images with significant interaction or avoidance determined by permutation test (*P <* 0.01). Rows representing the relationship of all other cell types surrounding a cell type of interest. Columns representing the relationship of a cell type of interest surrounding other cell types. Color in heatmap squares indicating the Spearman’s rank correlation of cell types across all IMC images in the nonresponders and responders. Highlighted interactions or avoidance (numbered black boxes) include: (1) lymphocytes; (2) MC4 cells and lymphocytes; (3) HLA-DR*^−^* myeloid cells and lymphocytes; (4) stromal SMA^+^ cells and immune cells. (c-g) Representative IMC images colored by marker and cell type showing the cell-cell interactions: B cells are surrounded by CD4 T cells (c), CD8 T cells are surrounded by CD4 T cells (d), CD8 T cells are surrounded by MC4 cells (e), CD4 T cells are surrounded by SMA^+^ stromal cells (f), CD8 T cells are surrounded by SMA^+^ stromal cells (g).

### Different TME archetypes based on multicellular compositions

We investigated how to translate the composition of single cells within TMEs into better stratification of melanoma to identify patients for immunotherapy. Using unsupervised hierarchical clustering on all the IMC images based the abundances of cell phenotypes that significantly differ in the IM regions of responders and non-responders, we obtained six TME archetypes that demonstrated distinct cell compositions, including three immune “hot” TMEs characterized by strong infiltration of CD4 T and B cells (H1), HLA-DR^+^CD11c^+^ myeloid derived cells (H2), and CD8 T cells (H3), respectively, and three immune “cold” TMEs with enrichment of CAIX^+^ tumor cells (C1), HLA-DR*^−^*CAIX^+^ myeloid derived cells (C2), and an archetype with no significant enrichment of any cell type (C3), respectively (Fig. 4a, HE and IMC images of example ROIs from each TME archetype shown in Fig. 4c). Signal pathway analysis with bulk RNA-seq data from paired samples also identified shared and distinct pathways of different TME archetypes (Fig. 4d and Supplementary Fig. S5). As expected, all immune “hot” TMEs showed multiple elevated signaling pathways that are correlated with adaptive and innate immune activation including IFN-*α*/*γ* response, allograft rejection, and complement pathway activities. H1 and H3 further showed unregulated inflammatory response and KRAS up-signaling pathways, while H2 was uniquely enriched for hallmarks of p53 pathway, and H3 uniquely enriched for hallmarks of apoptosis, IL2-STAT5 and IL6-JAK-STAT3 pathways. Immune “cold” TMEs were predominantly enriched for signaling pathways typically associated with cancer progression or immune evasion such as epithelial–mesenchymal transition and KRAS down signaling.

**Figure 4:**
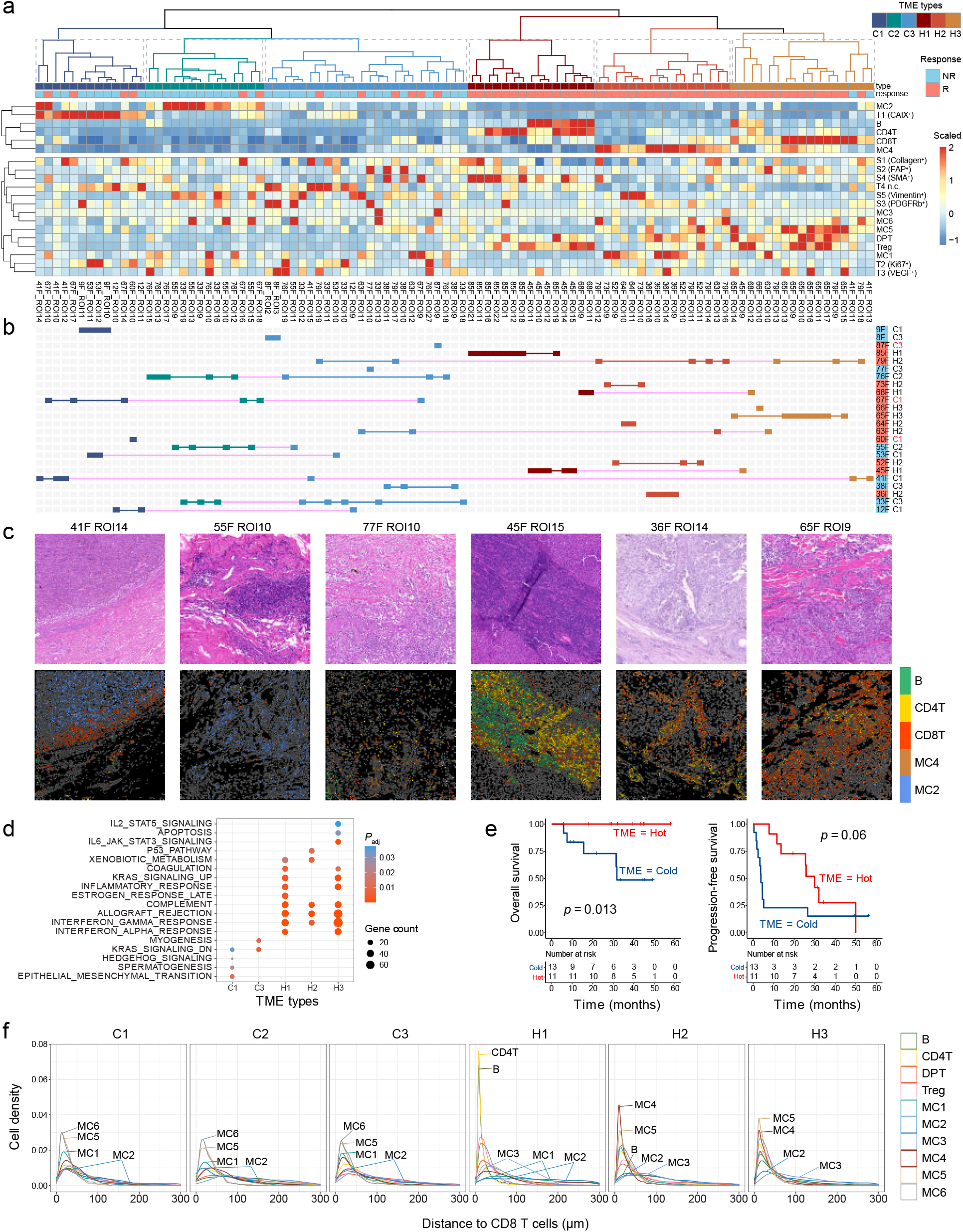
Identification of six distinct TME archetypes. (a) Heatmap showing scaled cell type abundance from the IM ROIs. Six TME archetypes are clustered by the level of selected cell types (MC2, Tumor CAIX^+^, B cells, CD4T, CD8T, MC4). (b) TME archetype patterns of each patient. If all ROIs from one patient are classified as having the same TME archetype, the patient is marked as the corresponding color of TME archetype. Patients who have ROIs that contain heterogeneous TME archetypes are indicated with magenta. (c) An example ROI from each TME archetype with HE image (top) and its corresponding IMC image (bottom) with cell phenotyping (B, CD4T, CD8T, MC2, MC4 cells). (d) Kaplan-Meier curves of OS (left) and PFS (right) for melanoma patients based on their TME archetypes. (e) Gene set enrichment analysis (GSEA) of genes with upregulated expression for each TME archetype. Significantly enriched gene sets (adjusted *P <*0.05, Benjamini–Hochberg method) from MSigDB HALLMARK collection are shown. (f) Histograms showing the nearest distance in µm between CD8 T cells and other immune/myeloid cells.

We further performed community analysis^24^ to investigate if single cells were organized differently in different TMEs. Using Louvain community detection^25^ to identify communities of multicellular units that were physically contacted with each other, followed by unsupervised clustering based on their cellular compositions using Phenograph, we obtained 19 common communities across all images (Methods, Supplementary Fig. S6a). Close examination on the community composition of different TME archetypes showed that each archetype had its own predominant multicellular communities (Supplementary Fig. S6b). For the immune “hot” TMEs, H1 was dominated by Community 3 that constituted large networks of CD4 T cells, B cells and CD8 T cells (Supplementary Fig. S7a); while in H2, the dominant community was Community 18 (Supplementary Fig. S7b) enriched for myeloid cells, primarily the HLA-DR^+^CD11c^+^ subtype MC4; and the majority community in H3 is Community 11 (Supplementary Fig. S7c) comprised of CD8 T cells that interacted with HLA-DR^+^ myeloid cells MC5. These multicellular communities were seldom found in immune “cold” TMEs. Instead, “cold” TME C1 contained the highest percentage of Community 6 (Supplementary Fig. S7d) that was characterized by CAIX^+^ tumor cells in close contact with MC2 and Collagen^+^ stromal cells; and C2 was mostly dominated by Community 4 (Supplementary Fig. S7e) enriched for networks of Vimentin^+^ stromal cells and HLA-DR*^−^*VEGF^+^ myeloid cells MC2. Finally, “cold” TME C3 showed a highly diffused cell distribution without any dominant communities.

We asked if the above classification of TMEs was associated with clinical outcome to anti-PD-1. Overall, by dividing the clustering results into “hot” and “cold” categories, this clustering achieved a classification accuracy of 79.3% (46 out of 58 responder ROIs classified as immune “hot”) for responders and 95.1% (39 out of 41 nonresponder ROIs classified as immune “cold”) for nonresponders on the ROI level (Fig. 4b). Analysis further revealed that ROIs from a same patient were in most cases highly homogeneous: most patients had ROIs from only one or two archetypes of the same immune “hot” or “cold” category (Fig. 4b). The exceptions included only two responders (79F, 63F) and one nonresponder (41F) who had ROIs from both immune “hot” and “cold” clusters. If we used majority voting to determine the TME archetype for each patient, all the 11 patients that were classified as immune “hot” were responders, representing an objective response rate (ORR) of 100%; and only 3 responders were from the immune “cold” patients, representing an ORR of 23.07%. Kaplan-Meier analysis revealed better overall survival (OS, *p* = 0.013) and progression-free survival (PFS, *p* = 0.06) in patients defined as immune “hot” (Fig. 4e).

Interestingly, despite the recognized important role of CD8 T infiltration to immunotherapy efficacy, only ROIs from H3 were characterized with significant CD8 T infiltration, representing only 6 out of 14 responders from this cohort. Close examination on different TMEs revealed highly different cell composition in the vicinity of CD8 T cells (Fig. 4f). In immune “hot” TMEs, the dominant cells surrounding CD8 T are either CD4 T and B cells in H1 or HLA-DR^+^ myeloid cells (MC4, MC5) in both H2 and H3, which is consistent with the recognized immune-enhancing actions governed by these cells. On the contrary, we observed significantly elevated accumulation of the HLA-DR*^−^* subtypes of myeloid cells (MC1, MC2) in close contact with CD8 T cells in all three immune “cold” TME archetypes (Fig. 4f), indicative of the potential role of these cells in creating an ICI resistant TME through mediating effector T cells functionality.

### Gene signature derived from distinct TME archetypes predicts anti-PD-1 therapy response

Recently, numerous gene expression signatures^26–29^ have been developed to study cellular composition of TMEs based on bulk RNA-seq data when single cell information is not available. Here, we investigated the consistency between our single cell analysis results from IMC data and the results from these signatures using RNA-seq data generated from adjacent serial section from the same samples in the PUCH cohort^30^. We performed correlation analysis between 29 curated GEP signatures^28^ and the cell type abundances estimated by averaging over all IMC ROIs for each sample (Supplementary Fig. S8a). Interestingly, we found an over-representation of CD8 T cells abundance in existing signatures despite that many of them have a putative target other than CD8 T cells. Among the 29 GEP signatures, the “Macrophages” signature shows the highest correlation with CD8 T cells abundance in the paired sample, followed by “Effector cells” and “T cells”. Other than CD8 T cells, DPT abundance showed strongest association with the “Effector cells” signature, while Treg abundance showed strongest association with the “Macrophage DC traffic” signature. Unfortunately, other than these three cell types, we did not find strong association between abundance of other cells and GEP signatures. For example, no surrogate GEP signatures for the abundances of myeloid subtypes were identified, while some myeloid cells (e.g., MC4), are strongly associated with clinical outcome to ICI in present study. We further performed correlation analysis between the cell type proportions estimated by the deconvolution method CIBERSORTx^29^ from bulk RNA-seq data and those estimated from IMC of the same sample, and similar observation can be made (Supplementary Fig. S8b). These findings suggested that existing RNA signature-and deconvolution-based methods for analyzing cellular compositions of TME could, at best, only capture the average cellular compositions of the whole tissue slide rather than their localized accumulations within TMEs due to the spatial heterogeneity of tummor tissues, while the latter are generally more essential for immunotherapy response prediction.

We further asked if it is possible to derive a global RNA-seq signature that could directly differentiate patients of different TMEs for immunotherapy outcome prediction without using cellular compositions as surrogates. To this end, we divided PUCH patients into immune “hot” and “cold” groups based on majority voting on their respective TMEs, and identified 20 significantly up-regulated immune-related genes and 4 significantly down-regulated immune-related genes in the immune “hot” group (Fig. 5a-b), where immune-related genes were defined as genes from the 770 curated cancer immune-related genes by Nanostring’s IO 360 panel (Methods). We then calculated a response score as the ratio of mean expressions of 20 up-regulated genes and 4 down-regulated genes to measure the anti-tumor immunity level for predicting anti-PD-1 outcome.

**Figure 5:**
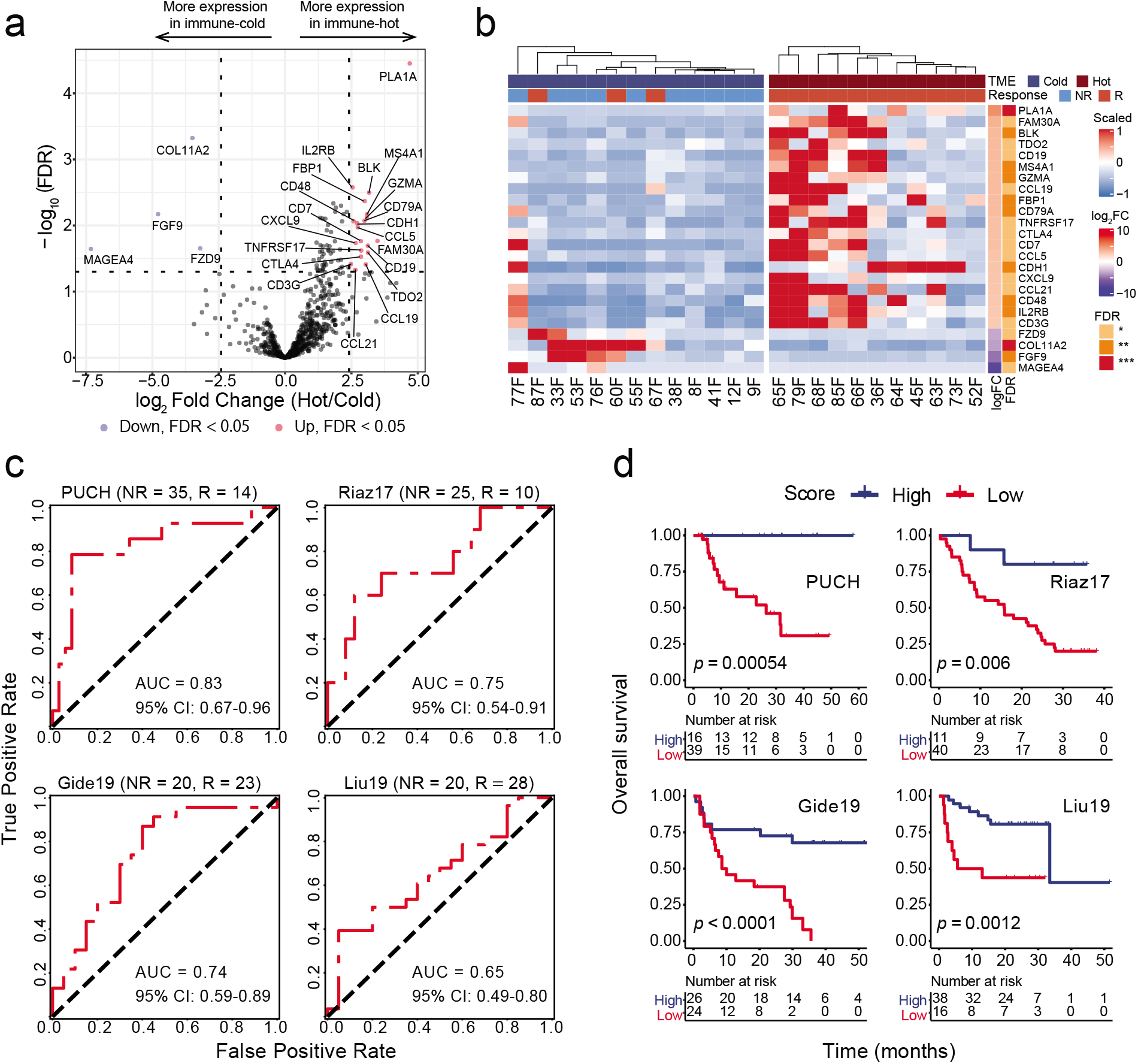
Prognostic impact of gene signature derived from TME archetypes. (a) Volcano plot showing the up-regulated genes (red) and down-regulated genes (blue) in patients with immune “hot” TME. (b) Heatmap depicting the expression of 24 DEGs (20 up-regulated genes, 4 down-regulated genes) from PUCH patients classified as immune “cold” and immune “hot” groups. (c) ROC curve of sensitivity vs 1-specificity of the prediction performance of prognostic score calculated by the 24 DEGs for PUCH dataset (discovery cohort) and other three public datasets (i.e., Riaz17, Gide19, Liu19). Patients with SD were not included in AUC calculation. (d) Kaplan-Meier curves of OS in melanoma patients with high versus low prognostic score calculated by the 24 DEGs for PUCH dataset (discovery cohort) and other three public datasets (i.e., Riaz17, Gide19, Liu19). Log-rank *p* values are shown.

To validate the performance of this signature, we analyzed RNA-seq data from the PUCH cohort and three independent external datasets from melanoma patients treated with anti-PD-1 (Riaz17^31^, Gide19^32^, Liu19^33^, Supplementary Table 3). Receiver operating characteristic (ROC) curve generated with clinical response data showed that the response score achieved an AUC of 0.83 (95% CI: 0.67-0.96) on PUCH, 0.75 (95% CI: 0.54-0.91) on Riaz17, 0.74 (95% CI: 0.59-0.89) on Gide19, and 0.65 (95% CI: 0.49-0.8) on Liu19, respectively (Fig. 5c). In addition, higher response scores were also associated with improved OS in the four datasets (Fig. 5d). Collectively, the above data demonstrated the potential value of using the response score derived from differentially expressed immune-related genes from patients of distinct TMEs as a biomarker for anti-PD-1 ICI treatments.

## Discussion

Our multidimensional interrogation of baseline melanoma tissue samples before anti-PD-1 treatment provided a systematic landscape of immune microenvironments of melanoma patients with different response to immunotherapy. Importantly, our results revealed highly heterogeneous TMEs from responders to immunotherapy, and only a subset of these TMEs have significant CD8^+^ T cells infiltration prior to immunotherapy, suggesting that anti-PD-1 therapy may have a much broader spectrum of mechanisms of action than only rejuvenating cytotoxic T cells that already reside in the TME. Indeed, rather than focusing on the specific state of a single cell type, a comprehensive recognition on the contributions from all cell types relevant to effective anti–PD-1 activity would be required for developing successful biomarkers in immunotherapy. For example, it is now well recognized that helper CD4^+^ T cells play a pivotal role in generating effective immune responses^34, 35^ and CD4^+^ T cell responses are required for optimal priming of antigen restricted CD8^+^ T cells and their maturation^36^. Although PD-1 is thought to predominantly restrain CD8^+^ effector T cells, recent studies show that its’ downstream effects further include activation of CD4^+^ T cells through targeting its costimulatory receptor CD28 by PD-1-recruited SHP2 phosphatase^37, 38^. Moreover, recent studies demonstrate that pre-existing T cells in TME have limited reinvigoration capacity^39^, and T cell responses to ICB are mainly derived from newly primed T cell clones from extrinsic repositories such as tumor-draining lymph nodes (TDLN)^40^, for which T cell priming through APCs that acquire tumor antigen and migrate to the TDLN would be required^41, 42^. For these reasons, as observed in the present study, enrichment of CD4^+^ T cell and/or myeloid derived APCs within TME could be a strong indicator to potential positive outcome to ICI in parallel to CD8^+^ T infiltration.

Tumors have been previously classified into immune “hot” with strong immune cell infiltrates or “cold” with sparse infiltration, and these pre-existing immune states are related to their potential responses to immunotherapy^28^. Our results supported this notion. Furthermore, empowered by multiplex single cell image analysis, we were able to identify multiple immune archetypes from both immune “hot” and “cold” TMEs. Each archetype is made up with a unique cellular community composition and charcterized by distinct dominant immune pathways, indicating that the previous TMEs delineations are incomplete to reveal the nuance of TMEs shaped by different tumor progression and immune evasion mechanisms. It is thus conceivable that such subdivision would enable us to further investigate the mechanisms behind different TME archetypes, from which better individualized therapeutic strategies based on archetypal assignments may be derived.

Although myeloid-derived cells are considered to associated with immune suppression within TME, it is now recognized that distinct myeloid cell subpopulations in the TME play different roles^43, 44^. Consistent with this notion, our results revealed two highly distinct archetypes of TMEs enriched for different myeloid cells. The first archetype (H2), of which the TMEs were all from responders, showed significant enrichment of HLA-DR^+^ myeloid cells, primarily the CD11c^+^ subtype MC4, but low pre-treatment lymphocytes infiltration, suggesting a potential seminal role played by this group of myeloid cells in mediating an inflammatory microenvironment towards positive outcome from anti-PD-1 treatment. The second archetype (C2), which was associated with poor clinical outcome to ICI, showed elevated accumulation of subtypes of highly exhausted myeloid cells with low HLA-DR expression and elevated VEGF and CAIX expressions (MC2), confirming their roles in immune suppression. Hence, in developing combination therapy that targets both T cell rejuvenation and macrophage depletion^45^, e.g., through CSF1R inhibitors^46^ combinning with ICB therapy, it may be necessary to identify the right target patients based on the composition of their myeloid infiltration as these inhibitors may lack the specificity to differentiate between protumor and antitumor myeloid cell subsets. In addition, repolarizing myeloid cells within TME to sustain or restore their tumoricidal activities through engaging pathogen recognition receptors (PRRs)^47^ or agonistic anti-CD40 antibody^48^ could be a promising combination therapeutic strategy to improve clinical response to ICI treatments for patients with this archetype of cold TME.

Other than TMEs enriched with exhausted myeloid cells, our results indicated the existence of another distinct immune “cold” TME archetype derived primarily from nonresponders (C1). TMEs of this archetype did not show strong infiltration of myeloid cells, but were characterized with enrichment of tumor cells with high expression of hypoxia signaling molecule CAIX. Hypoxic condition of tumor regions is typically arisen from increased oxygen consumption by rapidly proliferating tumor cells in combination with inadequate oxygen supply due to abnormal tumor angiogenesis^49^. Hypoxia-driven mechanisms allow tumor cells to continue to survive and proliferate in the hypoxic TME, while creating an inhospitable environment for immune cells through promoting apoptosis of T lymphocytes^50^ and DCs^51^, preventing effector T cells activation^52^ and their homing to the TME^50^, and promoting immune-suppressive stromal cells differentiation^53^, leading to tumor resistance to immunotherapy. Therefore, hypoxia may be exploited as a potential biomarker to identify this type of nonresponders, for whom strategies that combine methods to overcome hypoxia in cancer including hypxia-activated prodrugs (HAPs)^54^, inhibition of HIF signaling or its downstream pathways^55^, or supplemental oxygenation^53, 56^ with immunotherapy may be explored.

The limitations of this study include the small size of cohort and retrospective design. Nevertheless, our analysis has revealed highly heterogeneous multicellular features and their spatial interaction within a histological context of tumor TME, and confirmed that many of these features are associated with the clinical benefit of immunotherapy. Our results thus provide the basis for future studies on multicellular structures based on spatially resolved single-cell data for an in-depth characterisation of the tumor microenvironment, from which better methods to identify the right patients for different immunotherapy strategies can be derived. Moreover, our results further indicate that such knowledge is highly translatable, and can be exploited in multiple applications ranging from guiding the design of traditional bulk molecular tests for better patient segregation results despite their limitations in both spatial and single cell resolutions, or identification of targets for development of novel therapies.

## Methods

### Patient material

A total of 55 formalin-fixed, paraffin-embedded (FFPE) tumor tissue samples were obtained from melanoma patients with anti-PD-1 monotherapy at Peking University Cancer Hospital, Beijing, China. Patients were treated between March 2016 and March 2019 (Supplementary Table 1). Samples were collected from untreated patients before anti-PD-1 monotherapy.

Twenty-nine tissue samples were excluded as they did not meet the IMC experimental requirements, yielding the final cohort of 26 samples in the study (14 responders and 12 nonresponders). Clinical data, including sex, age, OS, PFS and clinical efficacy, were obtained from records of the patients with updated follow-up in Jun 2021. PFS was defined as the time from the date of treatment to disease progression or last contact. OS was defined as the time from treatment to death or last contact. Clinical efficacy to anti-PD-1 monotherapy was evaluated by Response Evaluation Criteria in Solid Tumors (RECIST) version 1.1^57^, including complete or partial response (CR/PR), stable disease (SD) and progressive disease (PD). All patients with CR/PR or SD were regarded as responders and PD patients are regards as nonresponders.

### Antibody conjugation and validation

An antibody panel of 35 proteins was designed to distinguish cell types and states, including immune, mesenchymal, proliferative and immune checkpoint proteins (Supplementary Table 2). Twenty-five labelled antibodies were purchased from Fluidigm (https://www.fluidigm.com) and the remaining 10 unlabelled antibodies were purchased from Abcam (https://www.abcam.com/). Antibodies from Abcam were conjugated with metals using Maxpar X8 Multimetal Labeling Kit (Fluidigm, 201300) following the manufacturer’s protocol. All conjugated antibody titration and specificity were tested by visual comparison of IMC images of some tissue slides from melanoma patients. Details about antibodies, metals and concentration used in the study can be found in Supplementary Table 2.

### Preparation and staining

Tissue slides were stained following IMC staining protocol (Fluidigm, PN400322) provided by Fluidigm. FFPE tumor samples were baked at 65% for 2h to remove all visible wax. Slides were deparaffinized in fresh xylene (10 min twice) followed by rehydration through a graded alcohol series (100%, 95%, 80%, 70% for 5 min each). Antigen retrieval was conducted in a 96°C water bath with Tris-EDTA buffer (pH9.0) for 30 min. Following cooling to 70°C at room temperature (RT), slides were then blocked with 3% BSA in PBS (Maxpar) for 45 min at RT in a hydration chamber. Meanwhile, the antibody cocktail was prepared in 0.5% BSA buffer mixed with the optimal dilution for each antibody (Supplementary Table 2). After blocking, slides were incubated with the antibody cocktail overnight at 4°C in a hydration chamber. The next day, each slide was washed twice with 0.2% Triton X-100 in PBS (Maxpar), and twice with PBS (Maxpar). For DNA staining, slides were incubated with Intercalator-Ir (Fluidigm, 201192A) in PBS (Maxpar) at RT for 30 min. Finally, slides were washed with deionized water twice and air-dried at least 20 min before IMC acquisition.

### Imaging mass cytometry

Images were acquired using a Hyperion Imaging System (Fluidigm). All operations were conducted following manufacturer’s procedure. Briefly, based on the HE-stained serial tissue section by a professional pathologist, we randomly selected ROIs at the CT or IM region. Images were laser ablated at 200 Hz, and raw data were acquired using a commercial acquisition software (Hyperion Imaging System, Fluidigm). The state of Hyperion Imaging System was monitored by interspersed acquisition of data from tuning slide (Fluidigm).

### IMC image processing, single-cell segmentation and quantification

We first checked the quality of every image by inspecting all marker staining patterns in the MCD Viewer (Fluidigm, v1.0.560.2). After quality control, a total of 158 images resulting in 662,266 single cells were used in the following analysis. Raw data (.mcd files) were converted to TIFF format using the imctools Python package (https://github.com/BodenmillerGroup/imctools). Then we used an in-house developed segmentation tool to perform single cell segmentation on each image^58^. The mean expression of 35 proteins of the segmented single cells were extracted using the “measure” module in scikit-image (Python package, v0.16.2) by overlaying the generated segmentation masks on the corresponding TIFF images. To improve the accuracy of cell protein expression value, all images for each channel were processed by our developed quantification pipeline^59^. Briefly, for each protein channel, a large number of random decoy cells were generated from IMC image regions that likely contained noise only. We then subtracted the mean expression of the decoy cells from those of the segmented single cells to remove the effect of the background noise on the quantification results. To remove the potential batch effect between ROIs, for each protein channel, we further identified positive cells by comparing the distribution of the expressions of the segmented single cells to that of the decoy cells with a false discovery rate (FDR) of 0.01, and normalized expressions of the segmented single cells across ROIs based on the expression of positive cells.

### Cell clustering analysis

Single cell protein expression data were clipped at the 99*^th^* percentile followed by min-max normalization. For cell type identification, 20 markers were used to define cell types: CD45, CD3, CD4, CD8a, FoxP3, CD20, CD68, CD14, CD16, CD11c, CD11b, IDO, Vimentin, *α*-SMA, E-cadherin, EpCAM, CAIX, VEGF, PDGFRb, Collagen I. Four main cell types (lymphoid cells, myeloid cells, stromal cells, tumor cells) were clustered and identified based on the protein expression pattern of each cluster. Then a second clustering was performed separately on all markers except for the immune checkpoint proteins and PD-L1, resulting in 20 distinguishable cell types from 77 clusters. All clustering analyses were performed with two consecutive steps: first meta-clusters were grouped with a self-organizing map implemented in FlowSOM^14^ (R package, v1.18.0), then Phenograph^15^ (R package, v0.99.1) was applied on the mean expression values of each group from FlowSOM clustering result to obtain the final clustering results. Cell type density was measured by the number of a certain cell type over total cells segmented from each image.

### Spatial analysis

To investigate cell-cell interactions, a permutation test method^23^ implemented in neighbouRhood (R package, v0.3.0) was used to determine whether the interactions/avoidances between or within cell types occurred more frequently than random observation. Briefly, cells were classified based on their protein expression values by cell clustering analysis as mentioned above, then a null distribution of cell interaction pairs was generated with 1,000 times permutation of random selection for each image. Observed interaction pairs were defined with a certain distance (distance between centroid of cells smaller than 20 µm). The *P*-value of interaction/avoidance between cell type A and B for each image was calculated as:

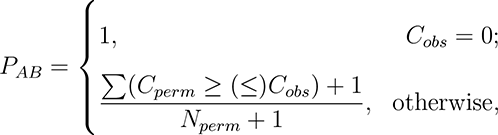

where *C_perm_* is the number of cell pairs (A, B) in each permutation, *C_obs_* is the actual number of cell pairs (A, B) given a defined distance, and *N_perm_* is the number of permutation. *P* values *≤* 0.01 were considered as significant interaction/avoidance between cell types. Interactions from observation and averaged interactions from permutation were compared by calculating log2 fold change (log2FC) to determine the extent of interactions/avoidance. Spatial proximity between two cell types were measured based on the distribution of the shortest distance from cells of one cell type to those of the other cell type on IMC images.

We further performed community analysis on IMC to identify common communities of multicellular units that existed across different TMEs^24^. Briefly, the IMC images were converted into topological neighbourhood graph in which cells were represented as nodes and direct cell-cell neighbouring pairs within 20 pixels (20 *µ*m between cell centroid) were represented as edges. Then we used the Louvain community detection method^25^ to identify highly interconnected spatial subunits in the graph. This analysis was performed on all cells to uncover the microenvironment communities across samples. Phenograph implemented in the cytofkit (R package, v0.99.0) was then used to identify recurring similar spatial cell type communities between samples based on minimum to maximum normalized percentages of cell types in each community.

### Measurement of inner-patient tumor heterogeneity

Each tumor sample represents a mixture of cells including lymphoid, myeloid, stroma and tumor cells. We used Shannon entropy (H) to characterize inner-patient heterogeneity based on annotated cell subtypes from cell clustering result. To account for the different number of cells per sample, we subsampled 1,000 cells from each sample *i* for three times and calculated its Shannon entropy of each occurred cell type frequency *P_c_* as:

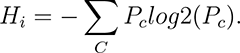

This analysis was performed on samples with different regions to investigate the cell type composition diversity in the CT or IM regions of patients using student t-test. We then compared the distribution of Shannon entropies of patients between responders and nonresponders.

### Identification of TME archetypes

We first selected the cell types that were differentially enriched between responders and nonresponders (log2FC *≥* 1.2, adjusted *p ≤* 0.05), and with a cell type density of at least 1% over total cells. The cell types that met these criteria were B, CD4 T, CD8 T, MC4, MC2, tumor (CAIX^+^) cells for ROIs in the IM, and MC2 and MC4 cells for ROIs in the CT. Hierarchical clustering was then conducted separately for ROIs in the IM on the basis of the Euclidean distance on the selected cell type abundances using hclust function with the Complete agglomeration method implemented in stats (R package, v3.6.3). For ROIs from IM, six distinct groups were generated by cutree function (R package stats) with k equal to 6. The resulting TME archetypes were further classified into two different categories (immune “hot”: H1, H2 and H3; immune “cold”: C1, C2 and C3) depending on their respective cell compositions.

### Whole-transcriptome RNA sequencing and external public datasets

Total RNA was extracted from unstained tissue slide which was adjacent serial section from the same FFPE tumor samples used in generating the IMC images. Sample RNA library construction and sequencing methods followed those as described in Cui *et al.*^30^. Briefly, RNA-seq reads were mapped by STAR^60^ and then quantified by RSEM^61^ to get fragments per kilobase of transcript per million mapped reads (FPKM) values at gene level. We further log2-transformed the read counts to avoid extremely skewed gene expression distributions.

In this study, we collected three RNA-seq datasets of melanoma patients treated with immunotherapy, together with their corresponding clinical information, including the Riaz17 (n = 51)^31^, Gide19 (n = 50)^32^, Liu19 (n = 54)^33^ datasets (Supplementary Table 3). We used the immunotherapy outcomes provided in the original papers following RECIST guidelines. For the Gide19 and Liu19 studies, only samples received anti-PD-1 monotherapy (nivolumab or pembrolizumab) were used. The RNA-seq raw data of Riaz17 and Gide19 datasets were obtained and processed by the same pipeline mentioned above to generate the gene expression data. For Liu19 datasets, the gene expression data were downloaded from respective references provided by the authors.

### Identification of DEGs, pathway analysis, and prognostic score calculation

Patients were classified into different TME archetypes based on majority voting, i.e., the archetype that had the most number of ROIs from a particular patient was assigned to the patient. Differential expression genes (DEGs) of each TME archetype were then identified using GLM function in edgeR (R package, v3.28.1) based on gene expressions of patients classified into that archetype vs. those of patients classified into archetypes of the opposite category. For example, DEGs of TME archetype H1 were derived based on gene expressions of patients from H1 vs. those from C1, C2 and C3. All DEGs with log2FC *≥* 1 and *p ≤* 0.05 for each TME archetype were inputted into ClusterProfiler (R package, v3.14.3) for gene set enrichment analysis on hallmark gene sets in Molecular Signatures Database (MSigDB v7.4).

To derive a prognostic gene signature, we identified DEGs between immune “hot” and immune “cold” patients. By using the common genes between DEGs and Nanostring’s IO 360 panel (770 curated cancer immune-related genes), we found 20 up-regulated genes (PLA1A, FAM30A, BLK, TDO2, CD19, MS4A1, GZMA, CCL19, FBP1, CD79A, TNFRSF17, CTLA4, CD7, CCL5, CDH1, CXCL9, CCL21, CD48, IL2RB, CD3G) and 4 down-regulated genes (MAGEA4, FGF9, COL11A2, FZD9). For each patient, the prognostic score was calculated as the ratio of mean expression of up-regulated genes to that of down-regulated genes.

### Deconvolution and ssGSEA score of 29 gene signatures from bulk RNA-seq data

Immune cell frequencies of bulk RNA-seq data were inferred using CIBERSORTx (https://cibersortx.stanford.edu/)^29^ which uses gene expression profile matrices from scRNA-seq data for deconvolution. We uploaded PUCH RNA-seq data, selected the absolute mode with online provided melanoma scRNA-seq data as the signature matrix, disabled quantile normalization and applied 100 permutations for deconvolution robustness.

Single-sample gene set enrichment analysis (ssGSEA, Python implementation by Bagaev et al.,^28^) was performed for 29 gene signatures which characterize four main TME groups (i.e., anti-tumor microenvironment, pro-tumor microenvironment, angiogenesis fibrosis, and malignant cell properties)^28^. To account for local region bias of IMC, the density of cell types for each sample were measured as the mean cell fraction of all ROIs taken from the same sample. Then we computed Spearman’s rank correlation and R-squared of linear regression model between cell type abundance from IMC and from RNA-seq data either by CIBERSORTx deconvolution or ssGSEA score of gene signature.

### Response prediction and survival analysis

To validate the prediction performance for each dataset, ROC curve was drawn based on the prognostic score using sklearn (Python package, v0.22.2). Kaplan-Meier analysis was performed to estimate OS or PFS using survival (R package, v3.2.3). For each dataset, we separated samples into two groups based on their prognostic scores with thresholds determined automatically by survminer (R package, v0.4.7). The log-rank test was used to assess the statistical comparison between two groups and a *p*-value *≤* 0.05 was considered significant. Univariable Cox proportional-harzards models adjusted by age were used to estimate the prognostic factors on survival, and the hazard ratio (HR) of each factor was reported using survival (R package, v3.2.3).

### Statistics and reproducibility

No statistical method was used to predetermine sample size and sample selection of this study was based on sample availability. All analyses were conducted using software R (version 3.6.3) and Python (version 3.7). The Wilcoxon rank-sum test was used for statistical analysis comparing continuous measurements, with Benjamini–Hochberg adjustment for all statistical tests involving multiple comparisons. An FDR-adjusted P *≤* 0.05 was considered significant. All box plots depict the median (the center line), interquartile range (IQR), and 1.5 times the IQR (whiskers), with outliers exceeding 1.5 times the IQR. For survival analysis, statistical significance between Kaplan-Meier curves were tested by the log-rank test. Correlation between cell type abundance was assessed by non-parametric Spearman’s rank correlation and the corresponding test. All statistical information used for experiments are defined in the figure legends.

## Code availability

All codes with data used to produce the results in this study are available at https://github.com/xmuyulab/IMC_melanoma.

## Ethics statement

The original studies were conducted in accordance with the Declaration of Helsinki and the International Conference on Harmonization Good Clinical Practice guidelines and approved by relevant regulatory and independent ethics committees from each study’s institution. All patients provided written informed consent before study entry.

## Authors contributions

W.Y., Y.K., and R.Y. supervised the research and designed the methodology. X.X., Q.G., C.C., Y.L., and M.W. performed image quantification, analyzed data and generated figures. L.Z and X.X. performed all IMC experiments. X.D. provided confirmatory pathology analyses. Y.K., Q.G., and C.C. provided patient samples and clinical input of the study. All authors assisted in writing the manuscript.

## References

1. Hodi, F. S. et al. Improved survival with ipilimumab in patients with metastatic melanoma. New England Journal of Medicine 363, 711–723 (2010).

2. Schachter, J. et al. Pembrolizumab versus ipilimumab for advanced melanoma: final overall survival results of a multicentre, randomised, open-label phase 3 study (KEYNOTE-006). The Lancet 390, 1853–1862 (2017). URL http://dx.doi.org/10.1016/S0140-6736(17)31601-X.

3. Robert, C. et al. Pembrolizumab versus ipilimumab in advanced melanoma. New England Journal of Medicine 372, 2521–2532 (2015).

4. Robert, C. et al. Pembrolizumab versus ipilimumab in advanced melanoma (KEYNOTE-006): post-hoc 5-year results from an open-label, multicentre, randomised, controlled, phase 3 study. The Lancet Oncology 20, 1239–1251 (2019). URL http://dx.doi.org/10.1016/S1470-2045(19)30388-2.

5. Farhood, B., Najafi, M. & Mortezaee, K. CD8+ cytotoxic T lymphocytes in cancer immunotherapy: A review. Journal of Cellular Physiology 234, 8509–8521 (2019).

6. Raskov, H., Orhan, A., Christensen, J. P. & Gögenur, I. Cytotoxic CD8+ T cells in cancer and cancer immunotherapy. British Journal of Cancer 124, 359–367 (2021).

7. Borst, J., Ahrends, T., Babala, N., Melief, C. J. & Kastenmüller, W. CD4+ T cell help in cancer immunology and immunotherapy. Nature Reviews Immunology 18, 635–647 (2018).

8. Richardson, J. R., Schöllhorn, A., Gouttefangeas, C. & Schuhmacher, J. CD4+ T cells: multitasking cells in the duty of cancer immunotherapy. Cancers 13, 596 (2021).

9. Sabado, R. L., Balan, S. & Bhardwaj, N. Dendritic cell-based immunotherapy. Cell Research 27, 74–95 (2017).

10. Liu, L. et al. Rejuvenation of tumour-specific T cells through bispecific antibodies targeting PD-L1 on dendritic cells. Nature Biomedical Engineering 1–13 (2021).

11. Helmink, B. A. et al. B cells and tertiary lymphoid structures promote immunotherapy response. Nature 577, 549–555 (2020). URL http://www.nature.com/articles/s41586-019-1922-8.

12. Petitprez, F. et al. B cells are associated with survival and immunotherapy response in sarcoma. Nature 577, 556–560 (2020).

13. Sun, X. et al. Tumour DDR1 promotes collagen fibre alignment to instigate immune exclusion. Nature 1–6 (2021).

14. Gassen, S. V., Callebaut, B., Helden, M., Lambrecht, B. N. & Saeys, Y. FlowSOM: Using self-organizing maps for visualization and interpretation of cytometry data. Cytometry Part A 87, 636–645 (2015).

15. Levine, J. H. et al. Data-driven phenotypic dissection of AML reveals progenitor-like cells that correlate with prognosis. Cell 162, 184–197 (2015).

16. Daud, A. I. et al. Tumor immune profiling predicts response to anti-PD-1 therapy in human melanoma. The Journal of Clinical Investigation 126, 3447–3452 (2016).

17. Hendriks, J. et al. CD27 is required for generation and long-term maintenance of T cell immunity. Nature Immunology 1, 433–440 (2000).

18. Menares, E. et al. Tissue-resident memory CD8+ T cells amplify anti-tumor immunity by triggering antigen spreading through dendritic cells. Nature communications 10, 1–12 (2019).

19. Han, J. et al. Resident and circulating memory t cells persist for years in melanoma patients with durable responses to immunotherapy. Nature Cancer 2, 300–311 (2021).

20. Wolf, Y., Anderson, A. C. & Kuchroo, V. K. TIM3 comes of age as an inhibitory receptor. Nature Reviews Immunology 20, 173–185 (2020).

21. Horton, B. L. et al. Lack of CD8+ T cell effector differentiation during priming mediates checkpoint blockade resistance in non-small cell lung cancer. Science Immunology 6, eabi8800 (2021).

22. Liu, Z. et al. Novel effector phenotype of Tim-3+ regulatory T cells leads to enhanced suppressive function in head and neck cancer patients. Clinical Cancer Research 24, 4529–4538 (2018).

23. Schapiro, D. et al. histoCAT: analysis of cell phenotypes and interactions in multiplex image cytometry data. Nature Methods 14, 873 (2017).

24. Jackson, H. W. et al. The single-cell pathology landscape of breast cancer. Nature 578, 615– 620 (2020).

25. Blondel, V. D., Guillaume, J.-L., Lambiotte, R. & Lefebvre, E. Fast unfolding of communities in large networks. Journal of statistical mechanics: theory and experiment 2008, P10008 (2008).

26. Lin, Y. et al. DAISM-DNN^XMBD^: Highly accurate cell type proportion estimation with in silico data augmentation and deep neural networks. bioRxiv 2020–03 (2021).

27. Aran, D., Hu, Z. & Butte, A. J. xCell: digitally portraying the tissue cellular heterogeneity landscape. Genome Biology 18, 220 (2017).

28. Bagaev, A. et al. Conserved pan-cancer microenvironment subtypes predict response to immunotherapy. Cancer Cell 39, 845–865 (2021).

29. Newman, A. M. et al. Determining cell type abundance and expression from bulk tissues with digital cytometry. Nature Biotechnology 37, 773–782 (2019).

30. Cui, C. et al. Ratio of the interferon-γ signature to the immunosuppression signature predicts anti-PD-1 therapy response in melanoma. NPJ Genomic Medicine 6, 1–12 (2021).

31. Riaz, N. et al. Tumor and microenvironment evolution during immunotherapy with nivolumab. Cell 171, 934–949 (2017).

32. Gide, T. N. et al. Distinct immune cell populations define response to anti-PD-1 monotherapy and anti-PD-1/anti-CTLA-4 combined therapy. Cancer Cell 35, 238–255.e6 (2019). URL https://doi.org/10.1016/j.ccell.2019.01.003.

33. Liu, D. et al. Integrative molecular and clinical modeling of clinical outcomes to PD1 blockade in patients with metastatic melanoma. Nature Medicine 25, 1916–1927 (2019).

34. Zhu, Z. et al. CD4+ T cell help selectively enhances high-avidity tumor antigen-specific CD8+ T cells. The Journal of Immunology 195, 3482–3489 (2015).

35. Bevan, M. J. Helping the CD8+ T-cell response. Nature Reviews Immunology 4, 595–602 (2004).

36. Alspach, E. et al. MHC-II neoantigens shape tumour immunity and response to immunotherapy. Nature 574, 696–701 (2019). URL http://www.nature.com/articles/s41586-019-1671-8.

37. Hui, E. et al. T cell costimulatory receptor CD28 is a primary target for PD-1-mediated inhibition. Science 355, 1428–1433 (2017).

38. Patsoukis, N., Wang, Q., Strauss, L. & Boussiotis, V. A. Revisiting the PD-1 pathway. Science Advances 6, eabd2712 (2020).

39. Yost, K. E. et al. Clonal replacement of tumor-specific T cells following PD-1 blockade. Nature Medicine 25, 1251–1259 (2019).

40. Dammeijer, F. et al. The PD-1/PD-L1-checkpoint restrains T cell immunity in tumor-draining lymph nodes. Cancer Cell 38, 685–700 (2020).

41. Topalian, S. L., Taube, J. M. & Pardoll, D. M. Neoadjuvant checkpoint blockade for cancer immunotherapy. Science 367 (2020).

42. Oh, S. A. et al. PD-L1 expression by dendritic cells is a key regulator of T-cell immunity in cancer. Nature Cancer 1, 681–691 (2020).

43. DeNardo, D. G. & Ruffell, B. Macrophages as regulators of tumour immunity and immunotherapy. Nature Reviews Immunology 19, 369–382 (2019). URL http://www.nature.com/articles/s41577-019-0127-6.

44. Neophytou, C. M. et al. The role of tumor-associated myeloid cells in modulating cancer therapy. Frontiers in Oncology 10, 899 (2020).

45. Mantovani, A., Marchesi, F., Malesci, A., Laghi, L. & Allavena, P. Tumour-associated macrophages as treatment targets in oncology. Nature Reviews Clinical Oncology 14, 399 (2017).

46. Cannarile, M. A. et al. Colony-stimulating factor 1 receptor (CSF1R) inhibitors in cancer therapy. Journal for immunotherapy of cancer 5, 1–13 (2017).

47. Rodell, C. B. et al. TLR7/8-agonist-loaded nanoparticles promote the polarization of tumour-associated macrophages to enhance cancer immunotherapy. Nature Biomedical Engineering 2, 578–588 (2018).

48. Schetters, S. T., et al. Monocyte-derived APCs are central to the response of PD1 checkpoint blockade and provide a therapeutic target for combination therapy. Journal for Immunotherapy of Cancer 8, e000588 (2020). URL https://jitc.bmj.com/lookup/doi/10.1136/jitc-2020-000588.

49. Noman, M. Z. et al. Improving cancer immunotherapy by targeting the hypoxic tumor microenvironment: new opportunities and challenges. Cells 8, 1083 (2019).

50. Mpekris, F. et al. Combining microenvironment normalization strategies to improve cancer immunotherapy. Proceedings of the National Academy of Sciences 117, 3728–3737 (2020).

51. Daniel, S., Sullivan, K., Labadie, K. & Pillarisetty, V. Hypoxia as a barrier to immunotherapy in pancreatic adenocarcinoma. Clinical and Translational Medicine 8, 1–17 (2019).

52. Damgaci, S. et al. Hypoxia and acidosis: immune suppressors and therapeutic targets. Immunology 154, 354–362 (2018).

53. Abou Khouzam, R., et al. Integrating tumor hypoxic stress in novel and more adaptable strategies for cancer immunotherapy. In Seminars in Cancer Biology, vol. 65, 140–154 (Elsevier, 2020).

54. Hunter, F. W., Wouters, B. G. & Wilson, W. R. Hypoxia-activated prodrugs: paths forward in the era of personalised medicine. British Journal of Cancer 114, 1071–1077 (2016).

55. Bhattarai, D., Xu, X. & Lee, K. Hypoxia-inducible factor-1 (HIF-1) inhibitors from the last decade (2007 to 2016): A “structure–activity relationship” perspective. Medicinal Research Reviews 38, 1404–1442 (2018).

56. Hatfield, S. M. et al. Immunological mechanisms of the antitumor effects of supplemental oxygenation. Science Translational Medicine 7, 277ra30–277ra30 (2015).

57. Schwartz, L. H. et al. RECIST 1.1 - Update and clarification: From the RECIST committee. European Journal of Cancer 62, 132–137 (2016).

58. Xiao, X., et al. Dice-XMBD: Deep learning-based cell segmentation for imaging mass cytometry. bioRxiv (2021).

59. Xiao, X., et al. IMCell^XMBD^: A statistical approach for robust cell identification and quantification from imaging mass cytometry images. bioRxiv (2021).

60. Dobin, A. et al. STAR: Ultrafast universal RNA-seq aligner. Bioinformatics 29, 15–21 (2013).

61. Li, B. & Dewey, C. N. RSEM: accurate transcript quantification from RNA-Seq data with or without a reference genome. Bioinformatics 12 (2011).

