## Supplementary for "Multiplexed imaging mass cytometry reveals distinct tumor-immune microenvironments linked to immunotherapy responses in melanoma"

Xiao et al

Table 1: Patient clinicopathologic characteristics.

| Characteristic | N = 26 |
|---|---|
| Sex, N (%) | |
| Male | 19 (73.1%) |
| Female | 7 (26.9%) |
| Age/Years, N (%) | |
| $\geq$ 60 | 9 (34.6%) |
| $\leq$ 60 | 17 (65.4%) |
| median (range) | 57 (30-72) |
| Race, N (%) | |
| Asian | 26 (100%) |
| Tumor site, N (%) | |
| Acral | 11 (42.3%) |
| Mucosal | 6 (23.1%) |
| Cutaneous | 5 (19.2%) |
| Unknown | 4 (15.4%) |
| Tumor thickness, N (%) | |
| $\leq$ 4 mm | 2 (7.7%) |
| >4 mm | 9 (34.6%) |
| Unknown | 15 (57.7%) |
| Ulceration | |
| With | 9 (34.6%) |
| Without | 4 (15.4%) |
| Unknown | 13 (50%) |
| TNM stage, N (%) | |
| I | 1 (3.8%) |
| II | 3 (11.5%) |
| III | 6 (23.1%) |
| IV | 16 (61.5%) |
| Efficacy, N (%) | |
| CR | 1 (3.8%) |
| PR | 10 (38.5%) |
| SD | 3 (11.5%) |
| PD | 12 (46.2%) |

Table 2: Antibodies and concentrations.

| Metal Tag | Targets | Clone | Dilution | Vendor |
|---|---|---|---|---|
| 141Pr | CD38 | EPR4106 | 1:100 | Fluidigm |
| 142Nd | PDGFRb | Y92 | 1:100 | Abcam (ab271835) |
| 143Nd | Vimentin | D21H3 | 1:100 | Fluidigm |
| 144Nd | CD14 | EPR3653 | 1:100 | Fluidigm |
| 145Nd | CD127 | EPR2955(2) | 1:100 | Abcam (ab240225) |
| 146Nd | CD16 | EPR16784 | 1:100 | Fluidigm |
| 147Sm | IDO | EPR20374 | 1:100 | Abcam (ab271990) |
| 148Nd | CD278 (ICOS) | D1K2T | 1:100 | Fluidigm |
| 149Sm | CD194 (CCR4) | L291H4 | 1:100 | Fluidigm |
| 150Nd | CD274 (PD-L1) | E1L3N | 1:100 | Fluidigm |
| 151Eu | CD134 (OX40) | Polyclonal | 1:100 | Fluidigm |
| 152Sm | CD45 | CD45-2B11 | 1:75 | Fluidigm |
| 153Eu | CD223 (LAG3) | D2G40 | 1:50 | Fluidigm |
| 154Sm | CD11c | Polyclonal | 1:100 | Fluidigm |
| 155Gd | FOXP3 | 236A/E7 | 1:100 | Fluidigm |
| 156Gd | CD4 | EPR6855 | 1:100 | Fluidigm |
| 158Gd | E-cadherin | 24E10 | 1:100 | Fluidigm |
| 159Tb | CD68 | KP1 | 1:200 | Fluidigm |
| 160Gd | EpCAM | EPR20532-222 | 1:50 | Abcam (ab232539) |
| 161Dy | CD20 | H1 | 1:100 | Fluidigm |
| 162Dy | CD8a | C8/144B | 1:100 | Fluidigm |
| 163Dy | VEGF | G153-694 | 1:75 | Fluidigm |
| 164Dy | CAIX | EPR23055-5 | 1:75 | Abcam (ab270401) |
| 165Ho | CD279 (PD-1) | EPR4877(2) | 1:100 | Fluidigm |
| 166Er | CD74 | LN2 | 1:100 | Fluidigm |
| 167Er | CD366 (TIM-3) | EPR22241 | 1:100 | Abcam (ab242080) |
| 168Er | Ki-67 | B56 | 1:100 | Fluidigm |
| 169Tm | Collagen type I | Polyclonal | 1:150 | Fluidigm |
| 170Er | CD3 | Polyclonal | 1:100 | Fluidigm |
| 171Yb | CD27 | EPR8569 | 1:75 | Fluidigm |
| 172Yb | FAP | Polyclonal | 1:100 | Abcam (ab53066) |
| 173Yb | CD11b | SP330 | 1:100 | Abcam (ab241408) |
| 174Yb | HLA-DR | LN3 | 1:100 | Fluidigm |
| 175Lu | Alpha-SMA | Polyclonal | 1:200 | Abcam (ab5694) |
| 176Yb | AXL | EPR19880 | 1:100 | Abcam (ab240396) |
| 191Ir | DNA1 | | | Fluidigm |
| 193Ir | DNA2 | | | Fluidigm |

Table 3: Cohorts used in this study.

| | PUCH [1] | Riaz17 [2] | Gide19[3] | Liu19 [4] |
|---|---|---|---|---|
| Cohort size, N | 55 | 51 | 50 | 54 |
| Nonresponse | 35 | 25 | 20 | 20 |
| Response | 14 | 10 | 23 | 28 |
| Stable disease | 6 | 16 | 7 | 6 |

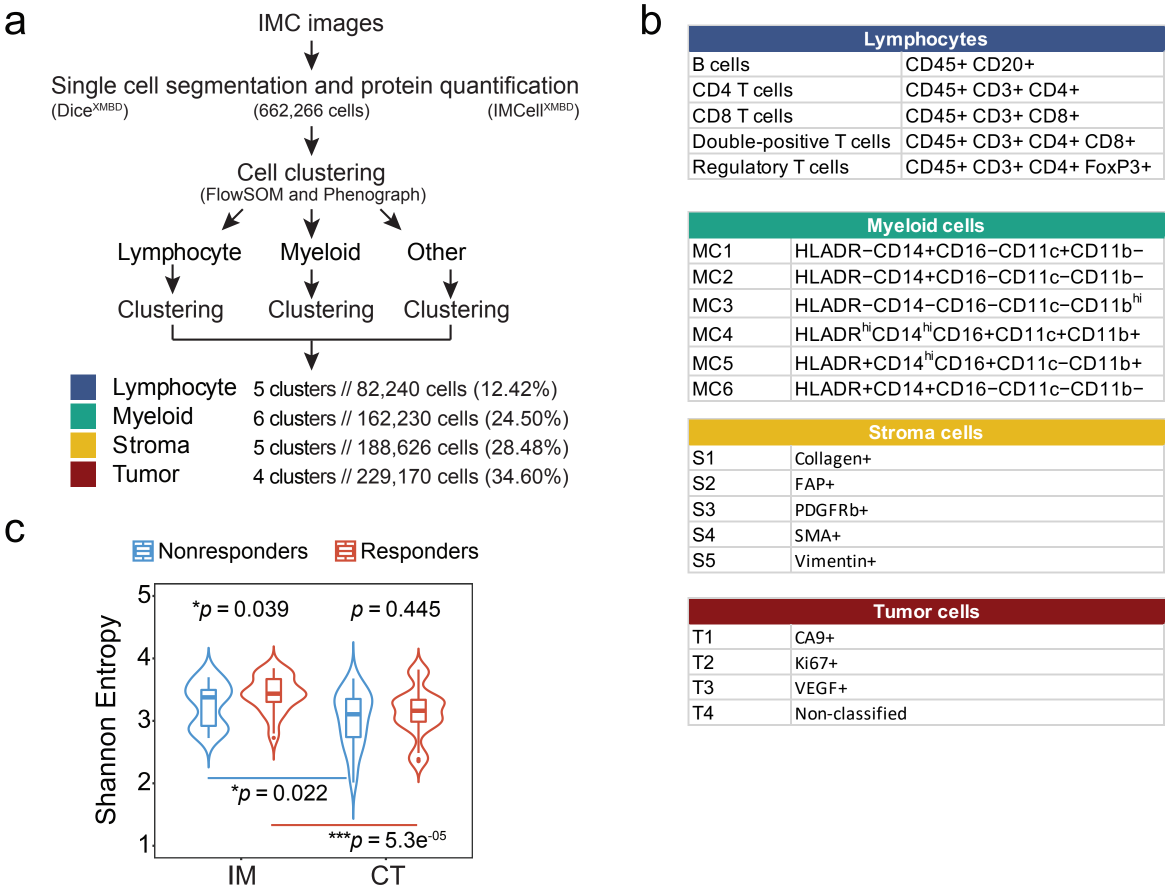


Figure 1: **The high-dimensional single cells phenotyping.** (a) Flowchart detailing the preprocessing and analytical steps taken at iterative subsets of the single-cell cohort. (b) Main cell types and subtypes with representing markers. (c) Inner-patient tumor heterogeneity measured by Shannon entropy for all samples (Wilcoxon rank sum test, $*p < 0.05$, $***p < 0.001$).

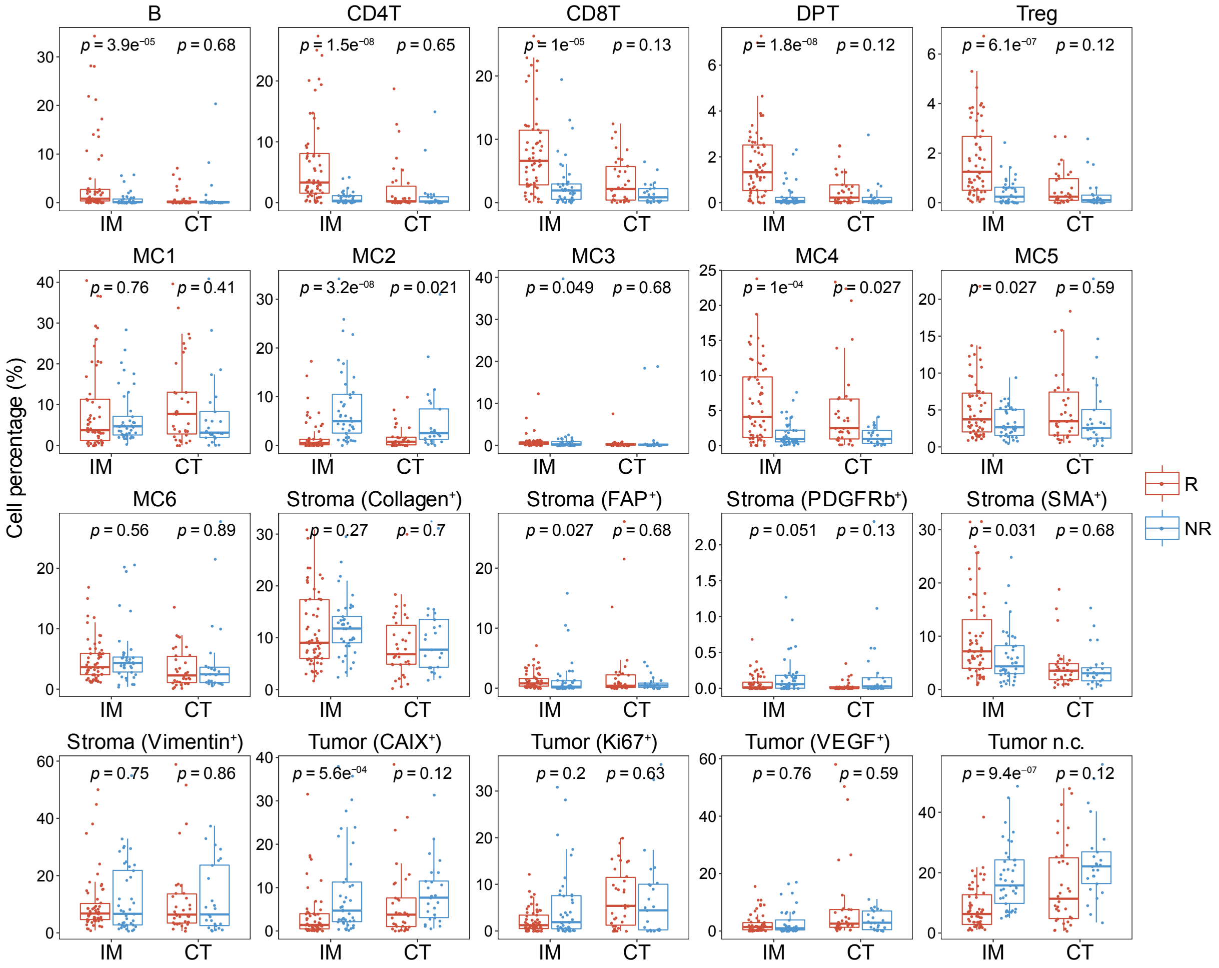


Figure 2: **The cell composition difference between responders and nonresponders.** Boxplots showing the proportion of cell type in ROIs from responders (R, red) and nonresponders (NR, blue). Each boxplot is shown with the median (the center line), interquartile range (IQR), and 1.5 times the IQR (whiskers), with outliers exceeding 1.5 times the IQR. Points in the boxplot represent the cell percentage of each IMC image (IM: n = 99, CT: n = 59). Comparisons were performed using Wilcoxon rank sum test and adjusted with Benjamini-Hochberg method.

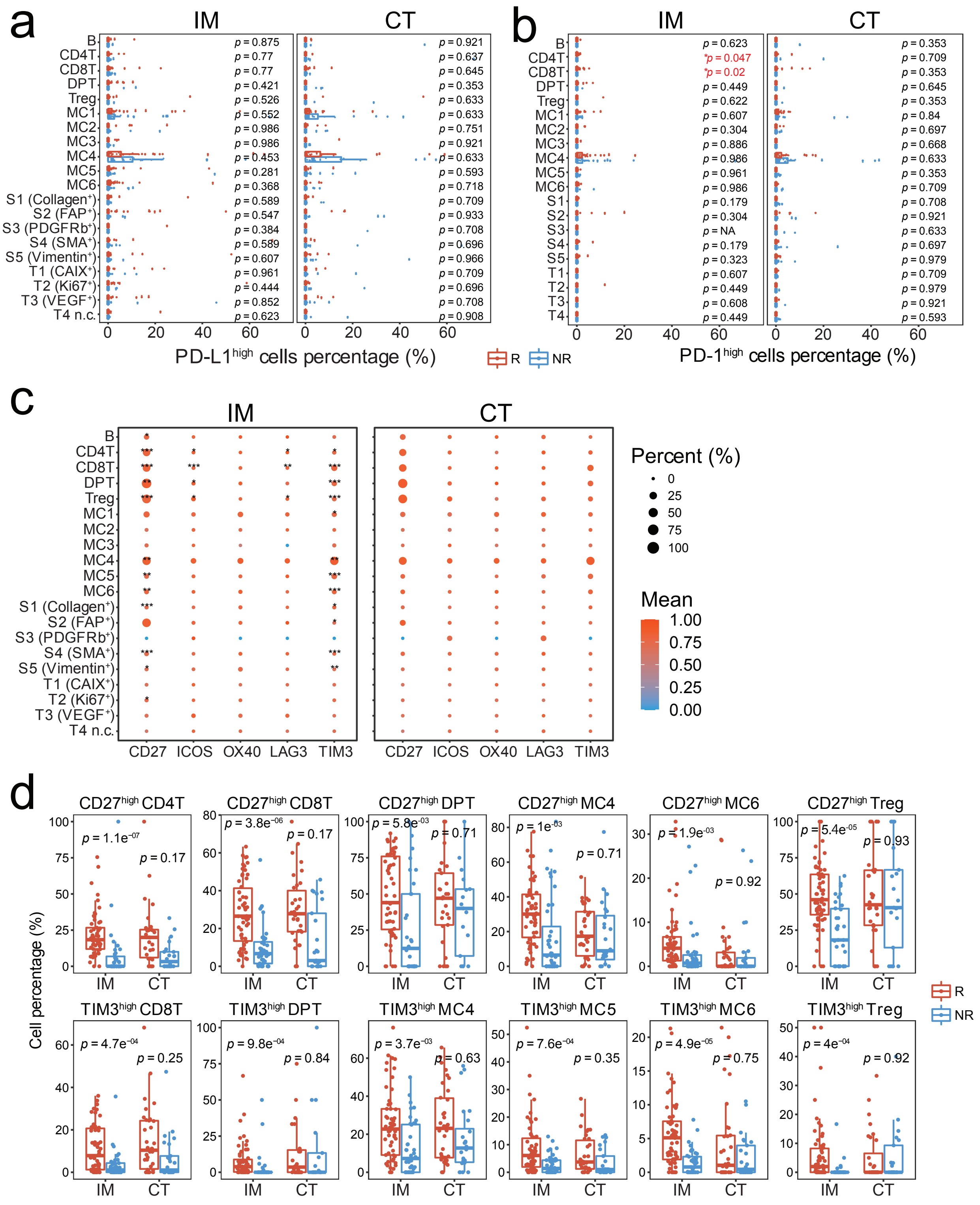
a
IM
CT
B
CD4T
CD8T
DPT
Treg
MC1
MC2
MC3
MC4
MC5
MC6
S1 (Collagen⁺)
S2 (FAP⁺)
S3 (PDGFRb⁺)
S4 (SMA⁺)
S5 (Vimentin⁺)
T1 (CAIX⁺)
T2 (Ki67⁺)
T3 (VEGF⁺)
T4 n.c.
p = 0.875
p = 0.77
p = 0.77
p = 0.421
p = 0.526
p = 0.552
p = 0.986
p = 0.986
p = 0.453
p = 0.281
p = 0.368
p = 0.589
p = 0.547
p = 0.384
p = 0.589
p = 0.607
p = 0.961
p = 0.444
p = 0.852
p = 0.623
p = 0.921
p = 0.637
p = 0.645
p = 0.353
p = 0.633
p = 0.633
p = 0.751
p = 0.921
p = 0.633
p = 0.593
p = 0.718
p = 0.709
p = 0.933
p = 0.708
p = 0.696
p = 0.966
p = 0.709
p = 0.696
p = 0.708
p = 0.908
0 20 40 60
PD-L1high cells percentage (%)
R
NR
b
IM
CT
B
CD4T
CD8T
DPT
Treg
MC1
MC2
MC3
MC4
MC5
MC6
S1
S2
S3
S4
S5
T1
T2
T3
T4
p = 0.623
*p = 0.047
*p = 0.02
p = 0.449
p = 0.622
p = 0.607
p = 0.304
p = 0.886
p = 0.986
p = 0.961
p = 0.986
p = 0.179
p = 0.304
p = NA
p = 0.179
p = 0.323
p = 0.607
p = 0.449
p = 0.608
p = 0.449
p = 0.353
p = 0.709
p = 0.353
p = 0.645
p = 0.353
p = 0.84
p = 0.697
p = 0.668
p = 0.633
p = 0.353
p = 0.709
p = 0.708
p = 0.921
p = 0.633
p = 0.697
p = 0.979
p = 0.709
p = 0.979
p = 0.921
p = 0.593
0 20 40 60
PD-1high cells percentage (%)
c
IM
CT
CD27 ICOS OX40 LAG3 TIM3
Percent (%)
0
25
50
75
100
Mean
1.00
0.75
0.50
0.25
0.00
d
CD27high CD4T
p = 1.1e-07
p = 0.17
CD27high CD8T
p = 3.8e-06
p = 0.17
CD27high DPT
p = 5.8e-03
p = 0.71
CD27high MC4
p = 1e-03
p = 0.71
CD27high MC6
p = 1.9e-03
p = 0.92
CD27high Treg
p = 5.4e-05
p = 0.93
TIM3high CD8T
p = 4.7e-04
p = 0.25
TIM3high DPT
p = 9.8e-04
p = 0.84
TIM3high MC4
p = 3.7e-03
p = 0.63
TIM3high MC5
p = 7.6e-04
p = 0.35
TIM3high MC6
p = 4.9e-05
p = 0.75
TIM3high Treg
p = 4e-04
p = 0.92
IM CT
Cell percentage (%)
R
NR

Figure 3: **The relationship between clinical response and expression level of checkpoint proteins per cell type.** The expression level was the relative abundance with respect to the total number of corresponding cell types. Positive cells were defined with expression values over three quarters of all cells. (a-b) Boxplots showing the relative percentage of PD-L1 (a) and PD-1 (b) positive cells in samples from responders (R, red) and nonresponders (NR, blue). (c) Dotplots displaying the relative proportion of positive cells using dot size, the mean expression level of selected proteins based on scaled expression value. (d) Boxplots showing the relative abundance of CD27/TIM3 positive cells in samples from responders (R, red) and nonresponders (NR, blue). Each boxplot is shown with the median (the center line), interquartile range (IQR), and 1.5 times the IQR (whiskers), with outliers exceeding 1.5 times the IQR. Points in the boxplot represent the cell percentage of each IMC image (IM: n = 99, CT: n = 59). Comparisons between responders and nonresponders were performed using Wilcoxon rank sum test and adjusted with Benjamini-Hochberg method. *$p < 0.05$, **$p < 0.01$, ***$p < 0.001$.

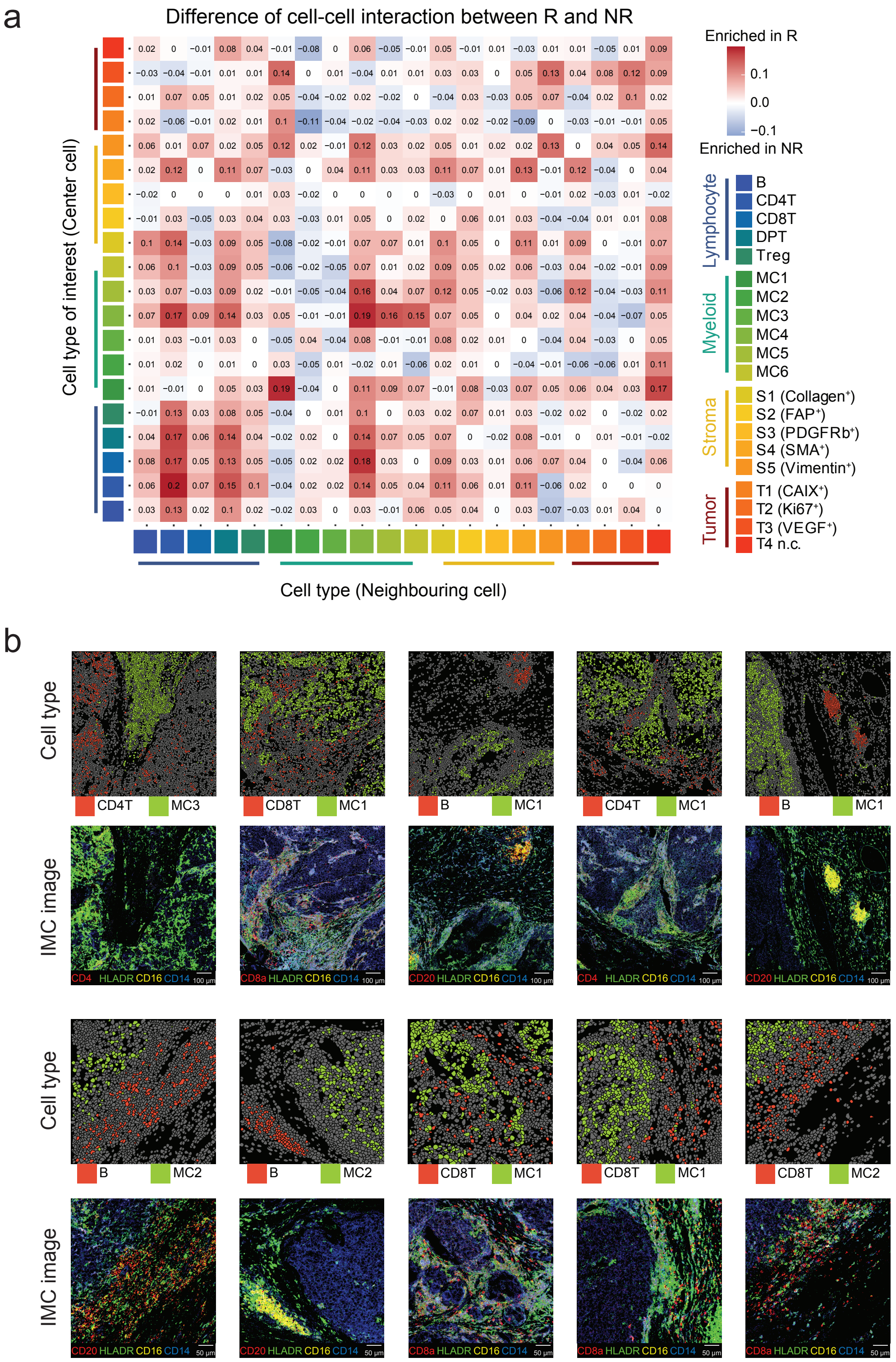


Figure 4: **Spatial analysis among cell phenotypes.** (a) Heatmap displaying frequencies of significant images determined by permutation test with $P < 0.01$ between responders and nonresponders. Red color indicating interactions that are more frequent in responders, and blue color indicating interactions that are more frequent in nonresponders. (b) Representative IMC images colored by cell type and marker showing cell-cell avoidance.

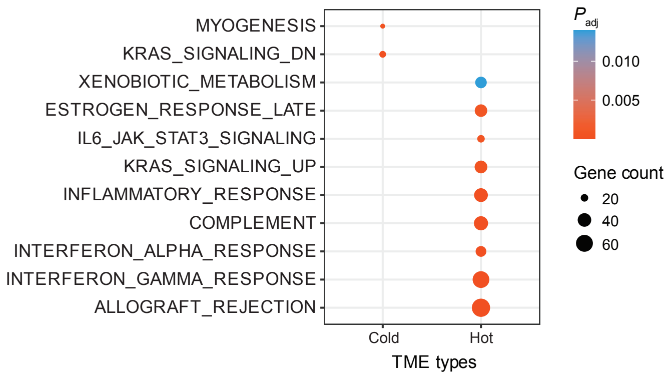


Figure 5: **Pathway analysis.** Gene set enrichment analysis (GSEA) of genes with up-regulated expression for immune "cold" and immune "hot" TME archetypes. Significantly enriched gene sets (adjusted $P < 0.05$, Benjamini-Hochberg method) from MSigDB HALLMARK collection are shown.

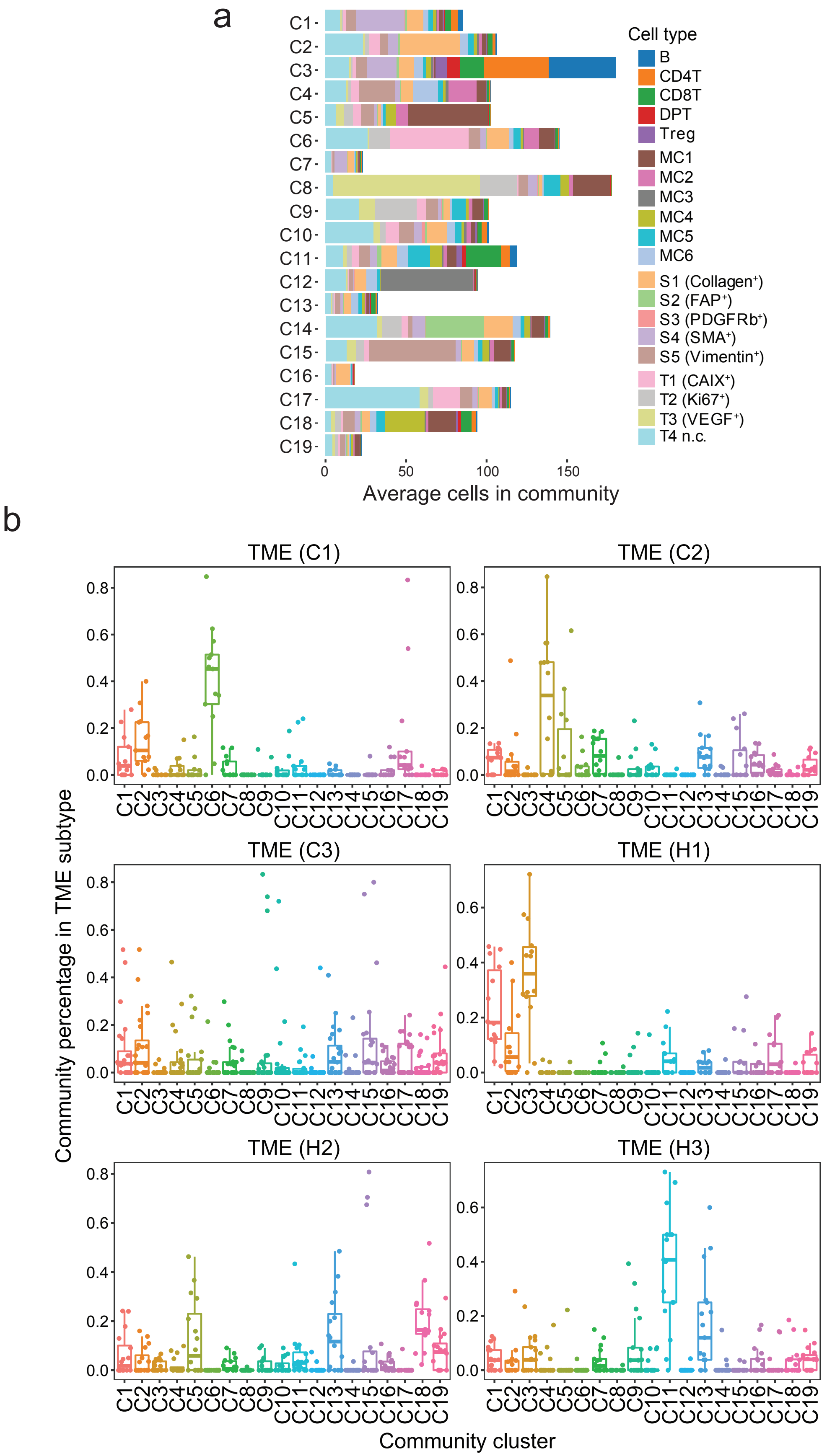
a
b
Cell type
B
CD4T
CD8T
DPT
Treg
MC1
MC2
MC3
MC4
MC5
MC6
S1 (Collagen⁺)
S2 (FAP⁺)
S3 (PDGFRb⁺)
S4 (SMA⁺)
S5 (Vimentin⁺)
T1 (CAIX⁺)
T2 (Ki67⁺)
T3 (VEGF⁺)
T4 n.c.
Average cells in community
TME (C1)
TME (C2)
TME (C3)
TME (H1)
TME (H2)
TME (H3)
Community percentage in TME subtype
Community cluster

Figure 6: **Community analysis.** (a) Microenvironment communities clustered by Phenograph based on the absolute percentage of cells per community, visualized on stacked bar plots indicating the average number of cells being assigned to each community. (b) Boxplots showing the percentage of community cluster in each TME archetype, colored by community cluster. Each boxplot is shown with the median (the center line), interquartile range (IQR), and 1.5 times the IQR (whiskers), with outliers exceeding 1.5 times the IQR. Points in the boxplot represent the community cluster percentage of each IMC image.

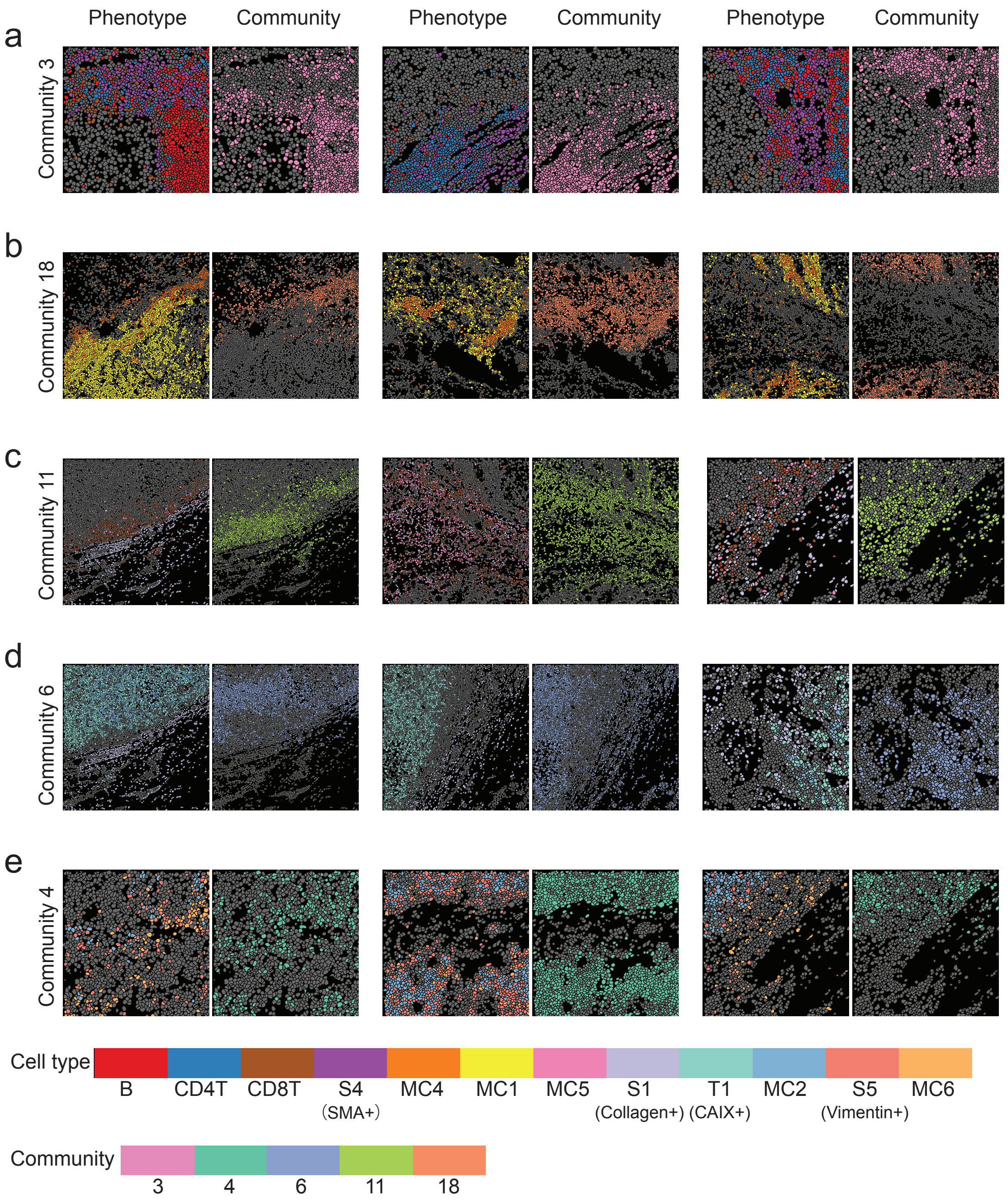


Figure 7: **Example pseudocolored images of communities.** (a-e) Rows showing three examples of community 3 (a), community 18 (b), community 11 (c), community 6 (d), and community 4 (e). Columns showing segmented cells colored by cell types (left) and community (right).

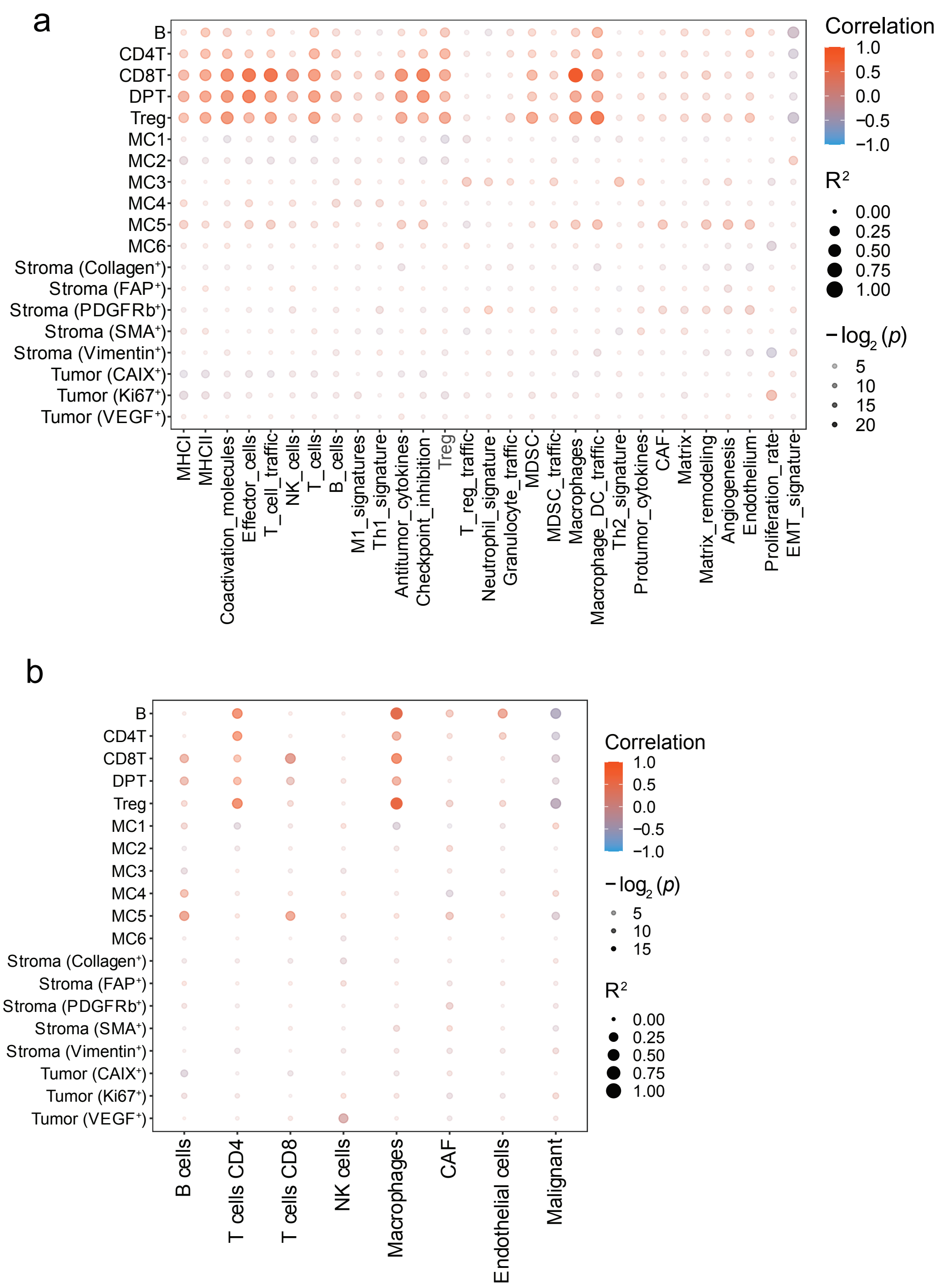


Figure 8: **The correlation between bulk RNA-seq data and IMC data.** (a-b) Spearman's rank correlation between the cell type abundances estimated by averaging over all IMC ROIs from a sample and estimated by ssGSEA score from 29 curated GEP signatures (a); and estimated by CIBERSORTx deconvolution method (b).

### References


1. Cui, C. *et al.* Ratio of the interferon-$\gamma$ signature to the immunosuppression signature predicts anti-PD-1 therapy response in melanoma. *NPJ Genomic Medicine* **6**, 1–12 (2021).

2. Riaz, N. *et al.* Tumor and microenvironment evolution during immunotherapy with nivolumab. *Cell* **171**, 934–949 (2017).

3. Gide, T. N. *et al.* Distinct immune cell populations define response to anti-PD-1 monotherapy and anti-PD-1/anti-CTLA-4 combined therapy. *Cancer Cell* **35**, 238–255.e6 (2019). URL https://doi.org/10.1016/j.ccell.2019.01.003.

4. Liu, D. *et al.* Integrative molecular and clinical modeling of clinical outcomes to PD1 blockade in patients with metastatic melanoma. *Nature Medicine* **25**, 1916–1927 (2019).
